# Disruption of CFAP418 interaction with lipids causes abnormal membrane-associated cellular processes in retinal degenerations

**DOI:** 10.1101/2022.06.13.495990

**Authors:** Anna M. Clark, Dongmei Yu, Grace Neiswanger, Daniel Zhu, J. Alan Maschek, Thomas Burgoyne, Jun Yang

**Author notes:** Corresponding Author: Jun Yang, John A. Moran Eye Center, University of Utah, 65 Mario Capecchi Drive, Bldg 523, Salt Lake City, Utah 84132. The authors have declared that no conflict of interest exists.

## Abstract

Syndromic ciliopathies and retinal degenerations are large heterogeneous groups of genetic diseases. *CFAP418* is a causative gene of both disorders, and its protein sequence is evolutionarily conserved. However, the pathogenic mechanism caused by *CFAP418* mutations is largely unknown. Here, we employed affinity purification coupled with mass spectrometry and quantitative lipidomic, proteomic, and phosphoproteomic approaches to address the molecular function of CFAP418 in mouse retinas. We showed that CFAP418 bound to lipid metabolism precursor phosphatidic acid (PA) and mitochondrion-specific lipid cardiolipin but did not form a tight and static complex with proteins. Loss of *Cfap418* led to membrane lipid imbalance and protein-membrane association alteration, which subsequently caused mitochondrial defects and membrane remodeling abnormalities in multiple vesicular trafficking pathways. Loss of *Cfap418* also increased the activity of PA-binding protein kinase Cα. Our results indicate that membrane lipid imbalance is a new pathological mechanism underlying syndromic ciliopathies and retinal degenerations, which is associated with other known causative genes for these diseases, such as RAB28 and BBS genes.

## Introduction

Eukaryotic cells have subcellular organelles that conduct specialized activities essential for cell growth, differentiation, homeostasis, survival, and function. Most organelles are separated from the cytoplasm by lipid bilayer membranes and exchange materials through intracellular trafficking to coordinate their activities and enable cells to act as an entity. During membrane intracellular trafficking, vesicles are budded at specific sites of donor organelles where cargos are sorted, travel through the cytoplasm, dock at, and fuse with the membranes of their target organelles. These processes are mediated by various membrane proteins and lipids (1, 2). The membrane lipids consist of numerous glycerophospholipids, sphingolipids, and sterols. The composition and distribution of these lipids determine the membrane’s biophysical and biochemical properties and associations with proteins. However, because of the significant complexity and diversity of membrane lipids and the scarcity of research tools, the mechanisms by which membrane lipids maintain cell organelle integrity and participate in membrane remodeling during vesicular transport have been understudied (3, 4).

Photoreceptors are highly polarized neurons and are an excellent cell model for studying membrane lipid homeostasis and intracellular vesicular trafficking (5, 6). From the apex to base, photoreceptors are composed of a specialized cilium known as the outer segment (OS), which is connected to the inner segment (IS), progressing through to the cell body within the outer nuclear layer (ONL), and finally the synaptic terminus within the outer plexiform (OPL) layer of the retina (Figure 1A). The IS is the cell compartment for most protein and membrane lipid biosynthesis. The synthesized proteins and membrane lipids are transported to other photoreceptor regions to generate different cell compartments during retinal development and to maintain cell homeostasis within mature retinas. The OS contains many tightly stacked membrane discs, where phototransduction occurs. These discs rapidly renew every 10 days in mammals to overcome the light-induced oxidative damage. Mutations in genes functioning in photoreceptor vesicular transport lead to photoreceptor cell death in a large heterogeneous group of inherited retinal degenerations (IRDs). The molecular basis of the intracellular vesicular transport in photoreceptors has not been well elucidated.

**Figure 1:**
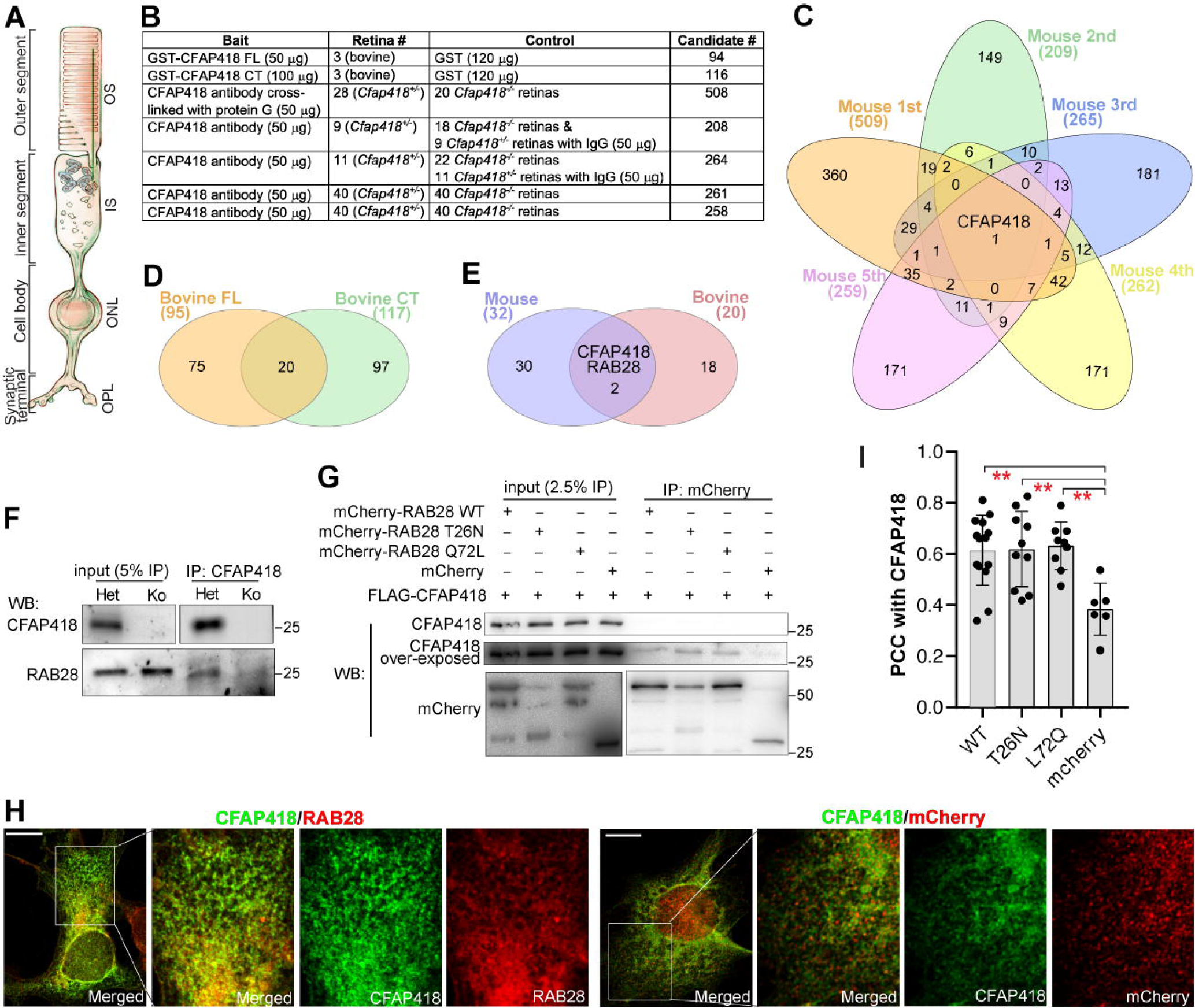
CFAP418 interacts weakly with RAB28 in retinas. **(A)** Schematic of photoreceptor subcellular compartments, which occupy the OS (outer segment), IS (inner segment), ONL (outer nuclear layer), and OPL (outer plexiform layer) in the retina. This schematic was adapted from (94). **(B)** Details of the conducted AP-MS experiments. **(C)** Overlap of proteins identified from mouse AP-MS experiments. **(D)** Overlap of proteins identified from bovine AP-MS experiments. **(E)** Overlap of potential CFAP418-interacting proteins from mouse and bovine retinas. **(F)** RAB28 was coimmunoprecipitated with CFAP418 from mouse retinas at 1 month of age. **(G)** mCherry-RAB28 proteins, but not mCherry, pulled down a small fraction of FLAG-CFAP418 in HEK293 cells. Note that FLAG-CFAP418 was only seen in over-exposed immunoblots. **(H)** Colocalization between FLAG-CFAP418 and mCherry-RAB28 but not mCherry. Framed regions are amplified and shown on the right. Scale bars: 10 μm. **(I)** PCCs of FLAG-CFAP418 with mCherry-RAB28 WT, T26N, L72Q, and mCherry alone. Data are represented as individual cells, mean, and standard error of the mean (SEM). **, *p* < 0.01 (Turkey’s multiple comparisons).

Cilia- and Flagella-Associated Protein 418 (*CFAP418*) is a causative gene for retinitis pigmentosa (RP), cone-rod dystrophy (CRD), and Bardet-Biedl syndrome (BBS) (7-15). While all the three IRDs affect photoreceptors, BBS is a syndromic ciliopathy and affects ciliated cells in multiple tissues. CFAP418 is 207 amino acids long in humans and has no known functional domains. Its sequence is highly conserved among species including the lower eukaryote *Chlamydomonas* (Figure S1A) and thus may play a fundamental role in eukaryotic cells. In zebrafish, *cfap418* knockdown leads to embryonic Kuppfer’s vesicle defects, retrograde melanosome transport delay, and visual impairment (7). We previously generated *Cfap418* knockout (*Cfap418^-/-^*) mouse models with different mutations (16). These *Cfap418^-/-^* mice display reduced electroretinogram responses, followed by photoreceptor cell death. *Cfap418^-/-^*photoreceptors exhibit extensive OS disc misalignment and OS membrane protein reduction. These phenotypes emerge at postnatal day 5 (P5), the onset of ciliogenesis, and become evident after P10 when OS grows robustly. CFAP418 is however localized to the photoreceptor IS. The exact function of CFAP418 remains unexplored.

Here, we applied unbiased omics approaches including affinity purification coupled with mass spectrometry (AP-MS) and quantitative mass spectrometry (MS) to investigate CFAP418-interacting proteins and the effects of *Cfap418* knockout on protein expression, phosphorylation, and membrane lipid composition in the retina. We unexpectedly discovered that CFAP418 binds directly to membrane lipid precursor phosphatidic acid (PA) and mitochondrion-specific lipid cardiolipin (CL), instead of forming a high-affinity and steady interaction with protein partners. Through these lipid bindings, CFAP418 maintains membrane lipid homeostasis, which is crucial for multiple membrane-associated cellular processes. This function of CFAP418 likely occurs in both ciliated and non-ciliated cells. Our study also reveals that disturbance of membrane lipid homeostasis is a novel pathological mechanism underlying ciliopathies and IRDs.

## Results

### Synthesis, degradation, and folding of OS membrane proteins are normal in *Cfap418^-/-^* photoreceptors

Rhodopsin is reduced in *Cfap418^-/-^*photoreceptors at P5 (16). We thus performed pulse and chase experiments to examine rhodopsin synthesis and degradation, respectively. Pulse labeling using [^35^S] methionine for up to 2 hours revealed no significant reduction of newly synthesized rhodopsin in *Cfap418^-/-^*retinas at P12 compared with a negative control cytoplasmic protein DPYSL2 (Figure S1B). Chase labeling using [^35^S] methionine in retinas for 2.5 hours, 3 hours, and overnight showed no evident changes in degradation of rhodopsin or transducin α subunit (GNAT1) at P8 and P10 (Figure S1C). Therefore, the rhodopsin synthesis and degradation rates appeared normal in *Cfap418^-/-^* retinas from P8 to P12, indicating that CFAP418 does not play an essential role in rhodopsin or other OS membrane protein synthesis or degradation in photoreceptors.

To test whether misfolding of OS membrane proteins occurred and induced ER stress and unfolded protein response (UPR) in *Cfap418^-/-^* retinas (17), we conducted RT-qPCR and found no changes in the mRNA expression of the three UPR pathway markers, ATF6, PERK, and IRE1α, in *Cfap418^-/-^*retinas at P15 and P30 (Figure S1D). We then tested whether ATF6 and IRE1α were activated using immunoblot analysis. We observed no cleavage of ATF6 into a 50-kDa fragment, which is the activated form of ATF6, and no elevation of phosphorylated IRE1α, the active form of IRE1α, in *Cfap418^-/-^*retinas at P16 and P30 (Figure S1E). Our data suggest that the OS membrane protein reduction in *Cfap418^-/-^* retinas is not due to protein misfolding.

### CFAP418 interacts weakly with RAB28 in the retina

We attempted to identify CFAP418-interacting proteins using AP-MS (Figure 1B and Table S1). In each of the five replicate experiments (Figure 1C), hundreds of proteins were coimmunoprecipitated with CFAP418 from *Cfap418^+/-^*retinas, but not with non-immunoglobin or from *Cfap418^-/-^* littermate retinas (negative controls). However, none of these proteins were coimmunoprecipitated with CFAP418 in all the five experiments, indicating that CFAP418 may not have a static or high-affinity binding to proteins. We also performed GST pulldown experiments from bovine retinal lysates using GST-tagged mouse CFAP418 full-length (FL) and C-terminal (CT) baits. The CT bait (169 – 209 aa, NP_080281), encoded by exon 6, is evolutionarily conserved in sequence and contains many pathogenic mutations identified in patients (Figure S1A) (7). Twenty proteins were pulled down by both CFAP418 FL and CT baits, but not by GST (Figure 1D). We compared these 20 proteins with the 32 proteins that were coimmunoprecipitated with CFAP418 at least three times from mouse retinas (Figure 1E). Only two proteins were shared between the two pools, CFAP418 and RAB28.

*CFAP418* (9, 15, 18) and *RAB28* (19-21) are both CRD genes. We thus investigated the potential interaction between CFAP418 and RAB28. Consistent with our AP-MS results, we detected RAB28 in the CFAP418 immunoprecipitate from mouse retinas in one of three coimmunoprecipitation experiments (Figure 1F). We then tested the direct interaction between RAB28 and CFAP418 and whether their interaction depended on the RAB28 GTP/GDP-binding status. We double-transfected HEK293 cells with FLAG-CFAP418 and mCherry-RAB28, RAB28_T26N (GDP-bound), or RAB28_Q72L (GTP-bound). mCherry-RAB28 proteins, but not mCherry (negative control), pulled down a small fraction of CFAP418 (∼0.1%), and GDP-RAB28 appeared to pull down more CFAP418 (Figure 1G) than other RAB28 proteins. CFAP418 and RAB28 are present in photoreceptor IS (16, 22). Because our CFAP418 antibodies could not detect the endogenous CFAP418 in photoreceptors by immunostaining and the volume of mouse photoreceptor IS is too small to examine detailed protein distribution, we examined the colocalization between CFAP418 and RAB28 in double-transfected COS-7 cells (Figures 1H and S3B). The Pearson correlation coefficients (PCCs) between CFAP418 and RAB28 WT, T26N, and L72Q were similar but significantly higher than the PCC between CFAP418 and mCherry (Figure 1I). The significant colocalization between CFAP418 and RAB28 was difficult to explain by their weak direction interaction, suggesting an indirect association between these two proteins in cells.

### CFAP418 binds to phosphatidic acid and cardiolipin in cell membranes

The weak interaction of CFAP418 with proteins prompted us to investigate the potential interactions of CFAP418 with membrane lipids. Both His- and GST-tagged mouse CFAP418 FL proteins were found to bind to PA and CL on membrane strips (Figures 2A and S2A). His-CFAP418 also bound weakly to lysophosphatidic acid (LPA) (Figure S2A). His- and GST-CFAP418 did not bind to phosphatidylcholine (PC), lysophosphatidylcholine (LPC), phosphatidylethanolamine (PE), phosphatidylserine (PS), phosphatidylglycerol (PG), phosphatidylinositol (PI), phosphoinositides (PIPs), triglyceride (TAG), diacylglycerol (DAG), cholesterol, sphingomyelin, sphingosine-1 phosphate, or sulfatide (Figures 2A and S2A).

**Figure 2:**
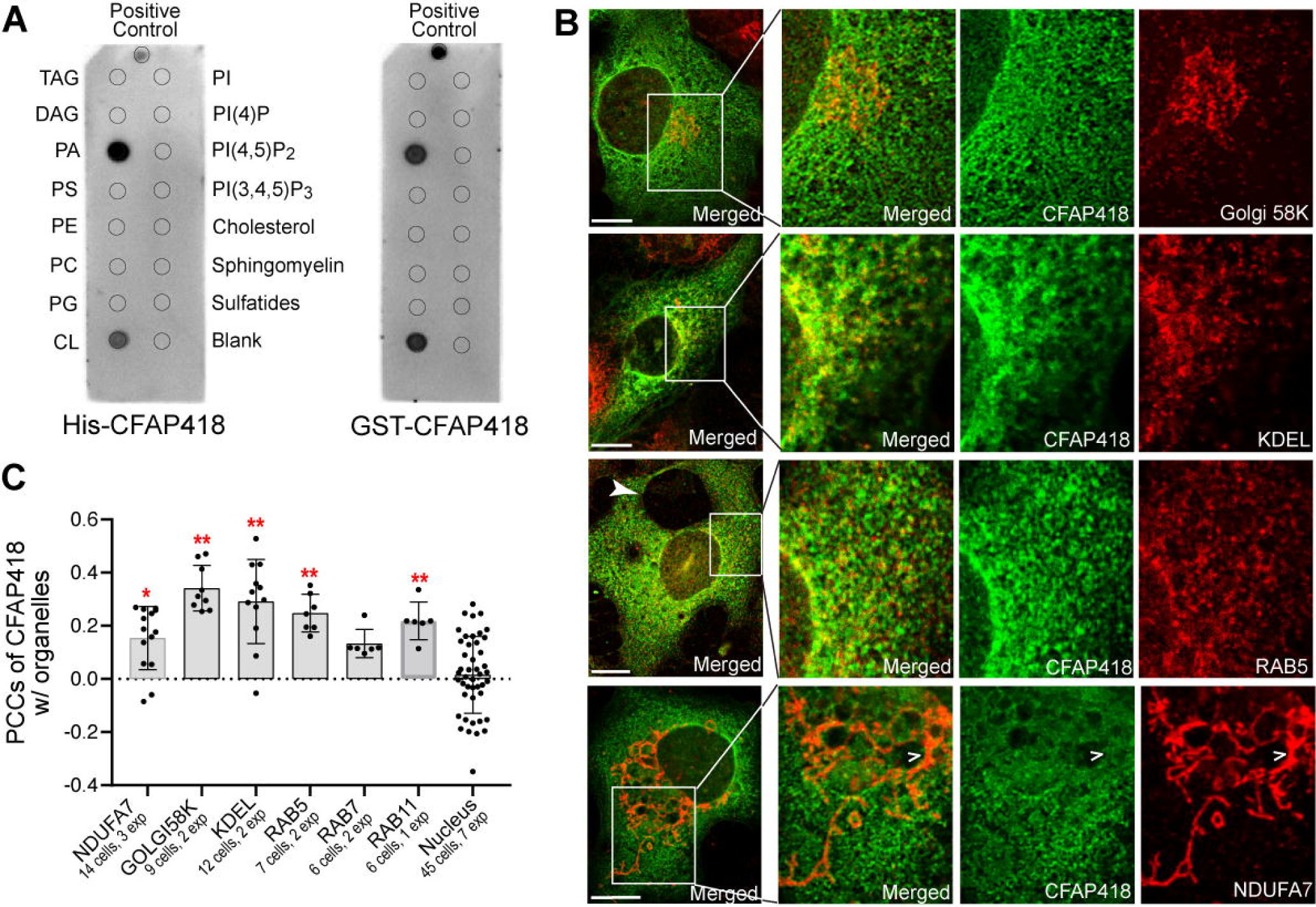
CFAP418 binds to PA and CL in various cell membranes. **(A)** His- and GST-CFAP418 proteins bind directly to PA and CL on membrane strips. The lipid arrangement is the same on the two membrane strips. Refer to the full lipid names in the Results section. **(B)** Representative COS-7 cells immunostained for FLAG-CFAP418 and various cell organelle markers. Filled arrowhead, large vacuole accumulation. Open arrowheads, localization of CFAP418 at mitochondrial edges. Scale bars: 10 μm. **(C)** PCCs of CFAP418 with different cell organelle markers. Nuclear dye Hoechst 33342 was used as a negative control. Data are represented as individual cells, mean, and SEM. * and **: *p* < 0.05 and 0.01, respectively (Turkey’s multiple comparisons with the nucleus group).

PA is virtually present in most cell membranes, and CL is exclusively found in mitochondrial membranes (3, 23). In our membrane association experiments, we observed a small amount of CFAP418 in the membrane fraction from *Cfap418*^+/-^ retinas (Figures 5G and 5H). Additionally, FLAG- and GFP-CFAP418 proteins appeared at the intracellular membranes and plasma membrane when transfected in COS-7 and HEK293 cells (Figures 2B, 5B, and not shown). To investigate which intracellular membranes CFAP418 associated with, we compared the distribution of transfected FLAG-CFAP418 with those of ER (KDEL), Golgi (Golgi 58K), mitochondrial (NDUFA7), and endosomal (RAB5, RAB7, and RAB11) markers in subconfluent COS-7 cells. The PCCs of CFAP418 with ER and Golgi apparatus were higher than those with endosomes and mitochondria (Figures 2B and 2C). Although the PCC between CFAP418 and mitochondrial marker NDUFA7 was relatively low, some CFAP418 signal contacted the edge of the NDUFA7 signal (open arrowhead in Figure 2B), implying the localization of CFAP418 at the mitochondrial outer membrane where ∼ 3% of CL exists (23, 24). As a control, the PCC of CFAP418 with a nuclear dye was around zero, consistent with our previous report that CFAP418 is absent in the nucleus (16). Together, our data demonstrate that CFAP418 binds to PA and CL in various cell membranes, including the ER, Golgi, endosome, and mitochondrial outer membranes.

### *Cfap418* deletion disrupts the membrane lipid composition in developing retinas

To examine whether and what membrane lipids were affected in *Cfap418*^-/-^ retinas, we performed a quantitative untargeted lipidomic analysis in the retinas of 16 *Cfap418^+/-^* and 16 *Cfap418^-/-^* littermates at P10. The data from one *Cfap418^-/-^* mouse was an outlier and was excluded (Figure S2B). Totally, 408 lipid species from 25 categories were detected (Table S2). The lipid composition in *Cfap418^+/-^* retinas was consistent with that previously reported (Figures S2C and S2D) (25). In *Cfap418^-/-^* retinas, the abundances of PC, lysophosphatidylserine (LPS), lysophosphatidylinositol (LPI), hexosylceramides (HexCer), and cholesterol were increased. In contrast, the abundances of EtherPC, EtherPE, acylglucuronosyldiacylglycerols (AcylGlcADG), CL, and ceramides (Cer) were reduced (Figure 3A). Cholesterol was the most increased with a ratio of *Cfap418^-/-^*to *Cfap418^+/-^* abundance (FC) at 1.26 ± 0.06 (mean ± SEM, *p* = 0.00086) and CL, as a group of CFAP418-binding lipids, was the second most reduced lipid category (FC = 0.75 ± 0.03, *p* = 0.0010). Within the CL category, CL68:4, CL68:5, CL70:4, and CL72:5 were significantly reduced (FC < 0.95 and False Discovery Rate [FDR] q < 0.05, Figure 3C and Table S3). Note that the X:Y in the lipid nomenclature indicates the numbers of carbons and double bonds, respectively, in the lipid acyl chains. Consistently, Metabolite Set Enrichment Analysis (MSEA) on the quantitative lipidomic profiles without any cutoff thresholds found that the lipid changes were enriched in the CL (FDR = 0.0117), cholesterol (FDR = 0.0117), and PC (FDR = 0.0117) categories in *Cfap418^-/-^* retinas.

**Figure 3:**
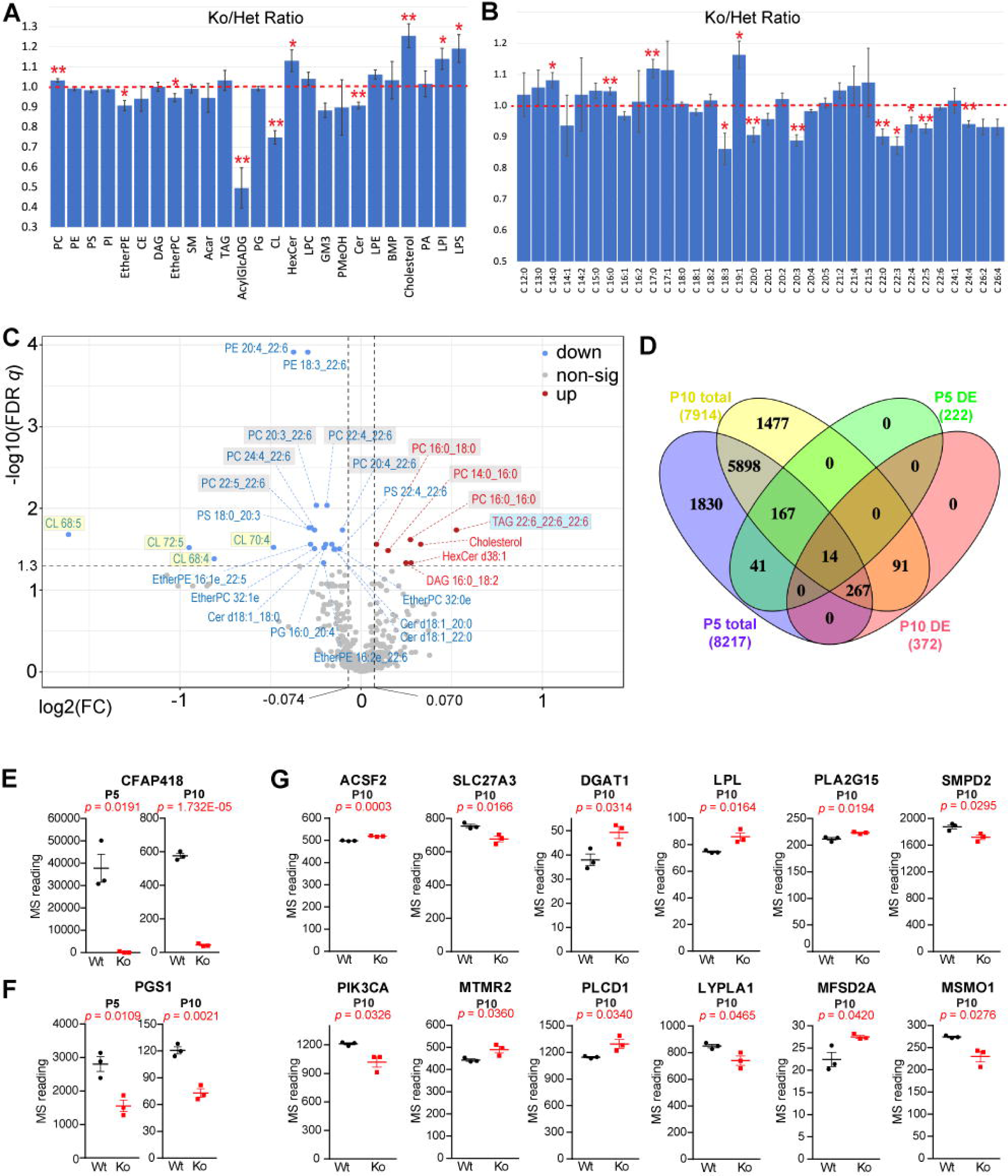
Abnormal membrane lipid composition and lipid regulated protein expression in developing *Cfap418^-/-^* retinas. **(A)** Affected membrane lipid categories in P10 *Cfap418^-/-^*retinas. **(B)** Affected acyl chains in P10 *Cfap418^-/-^* retinal membrane lipids. **(C)** Volcano plot showing fold changes of individual lipid species between P10 *Cfap418^+/-^* and *Cfap418^-/-^* retinas. **(D)** Overlap of the total detected proteins and DE proteins between the P5 and P10 proteomes. **(E)** Scarcity of CFAP418 in *Cfap418^-/-^* retinas validates our quantitative proteomic data. **(F)** Reduced PGS1 expression in P5 and P10 *Cfap418^-/-^* retinas. **(G)** Altered expressions of membrane lipid metabolic enzymes and transporters in P10 *Cfap418^-/-^* retinas. Bar charts: mean ± SEM; * and **, *p* < 0.01 and 0.05, respectively (Student’s t-test). Dot plots: data of individual mice and mean ± SEM.

Further analysis revealed that the acyl chains with fewer carbons and zero or one double bond were increased (i.e., C14:0, C16:0, C17:0, and C19:1), and the acyl chains with more carbons and more double bonds were reduced (i.e., C18:3, C20:0, C20:3, C22:0, C22:3, C22:4, C22:5, and C24:4) in *Cfap418^-/-^* membrane lipids (Figure 3B). This phenotype was evident in the PC species that were altered in *Cfap418^-/-^* retinas (Figures 3C and S2E). PC14:0_16:0, PC16:0_16:0, and PC16:0_18:0 were increased, while PC20:4_22:6, PC22:4_22:6, PC22:5_22:6, and PC24:4_22:6 were decreased. In addition to the affected CL, PC, and cholesterol species, 6 other glycerophospholipids, 4 ether glycerophospholipids, and 3 sphingolipids were reduced, and 2 glycerolipids and 1 sphingolipid were increased in *Cfap418^-/-^* retinas (Figure 3C and Table S3). Because of the low abundance of PA in membrane lipids (3, 26), we detected only one PA species, which was unaffected in *Cfap418^-/-^* retinas.

### Lipid metabolic enzyme and transporter expressions are altered in developing *Cfap418*^-/-^ retinas

We surveyed the protein expression defects in *Cfap418^-/-^*retinas in an unbiased manner using proteome-wide quantitative MS in three pairs of wild-type and *Cfap418^-/-^* littermates at each time point. At P5, label-free quantitative MS detected 193 down-regulated and 29 up-regulated proteins (*p* < 0.05, Table S3) from a total of 8,218 detected proteins in *Cfap418^-/-^* retinas (Figure 3D and Table S2). At P10, tandem mass tag (TMT)-labeling quantitative MS identified 237 down-regulated and 146 up-regulated proteins (*p* < 0.05, Table S3) from a total of 7,914 detected proteins in *Cfap418^-/-^*retinas (Figure 3D and Table S2). As expected, *Cfap418* knockout led to barely detectable CFAP418 protein expression in both P5 and P10 retinas (Figure 3E).

Among the 333 known membrane lipid metabolic enzymes and transporters (4), 186 were detected in the P5 and P10 retinal proteomes. Phosphatidylglycerophosphate synthase (PGS1), the enzyme responsible for the first step of CL synthesis in mitochondria (27, 28), was reduced by ∼45% and 40% in P5 and P10 *Cfap418^-/-^*retinas, respectively (Figure 3F), which was consistent with our observed CL reduction in *Cfap418^-/-^*retinas. No other membrane lipid metabolic enzymes or transporters showed changes in P5 *Cfap418^-/-^* retinas. However, at P10, 11 more membrane lipid metabolic enzymes and transporters were differentially expressed (DE, Figure 3G). Some of these changes were consistent with the membrane lipid changes we observed (Figure 3). For example, ACSF2 and SLC27A3 are medium-chain and very long-chain acyl-CoA synthetases in mitochondria, respectively (29, 30). They function in the de novo synthesis and remodeling of acyl chains in glycerophospholipids (4, 31). The ACSF2 increase and the SLC27A3 reduction may lead to the opposite changes of short and long acyl chains in *Cfap418^-/-^* glycerophospholipids. PLA2G15 functions as both phospholipase A2 and ceramide acyltransferase toward PC, PE, PS, and PG (32, 33). Its increase may explain the LPS and LPI increases and the subsequent acyl chain remodeling. The increases of TAG transporter LPL (34) and TAG synthetase DGAT1 (35, 36) may contribute to the increase of TAG 22:6_22:6_22:6 in *Cfap418^-/-^*retinas. The reduction of SMPD2, a sphingomyelinase for ceramide generation (37), may be responsible for the ceramide reduction in *Cfap418^-/-^* retinas. Furthermore, MTMR2, PLCD1, and PIK3CA participate in PIP metabolism and signaling (38-41). Their alterations may affect the PIP abundances in *Cfap418^-/-^* retinas, which were not detected in our lipidomic studies due to the extremely low PIP abundance in retinal tissues (42). Because most DE lipid metabolic enzymes and transporters were affected in P10 but not P5, their expression changes are probably secondary to the disrupted membrane lipid composition in *Cfap418^-/-^* retinas.

### Mitochondrial morphology and function are abnormal in developing *Cfap418^-/-^* photoreceptors

Considering the binding of CFAP418 with CL, significant reductions of CL and CL synthetase PGS1 in *Cfap418^-/-^* retinas, and the specific localization of CL in mitochondria, we examined the mitochondrial morphology and structure in *Cfap418^-/-^* photoreceptors using transmission electron microscopy (TEM). At P10, the mitochondria showed a dynamic polymorphous shape in photoreceptors, which seemed slightly more irregular in *Cfap418^-/-^* photoreceptors than in *Cfap418^+/-^* photoreceptors. This phenotype became obvious in mature photoreceptors at P21, P28, and P60. At these time points, most mitochondria had a straight long cylindrical shape and were located immediately parallel to the plasma membrane of *Cfap418^+/-^*photoreceptor IS (Figure 4A). In *Cfap418^-/-^* littermate photoreceptors, the mitochondria were located normally, and the size and number of the cristae within mitochondria appeared normal. However, the mitochondria displayed an irregular cylindrical shape with bumps and constrictions along their longitudinal axis (Figure 4A).

**Figure 4:**
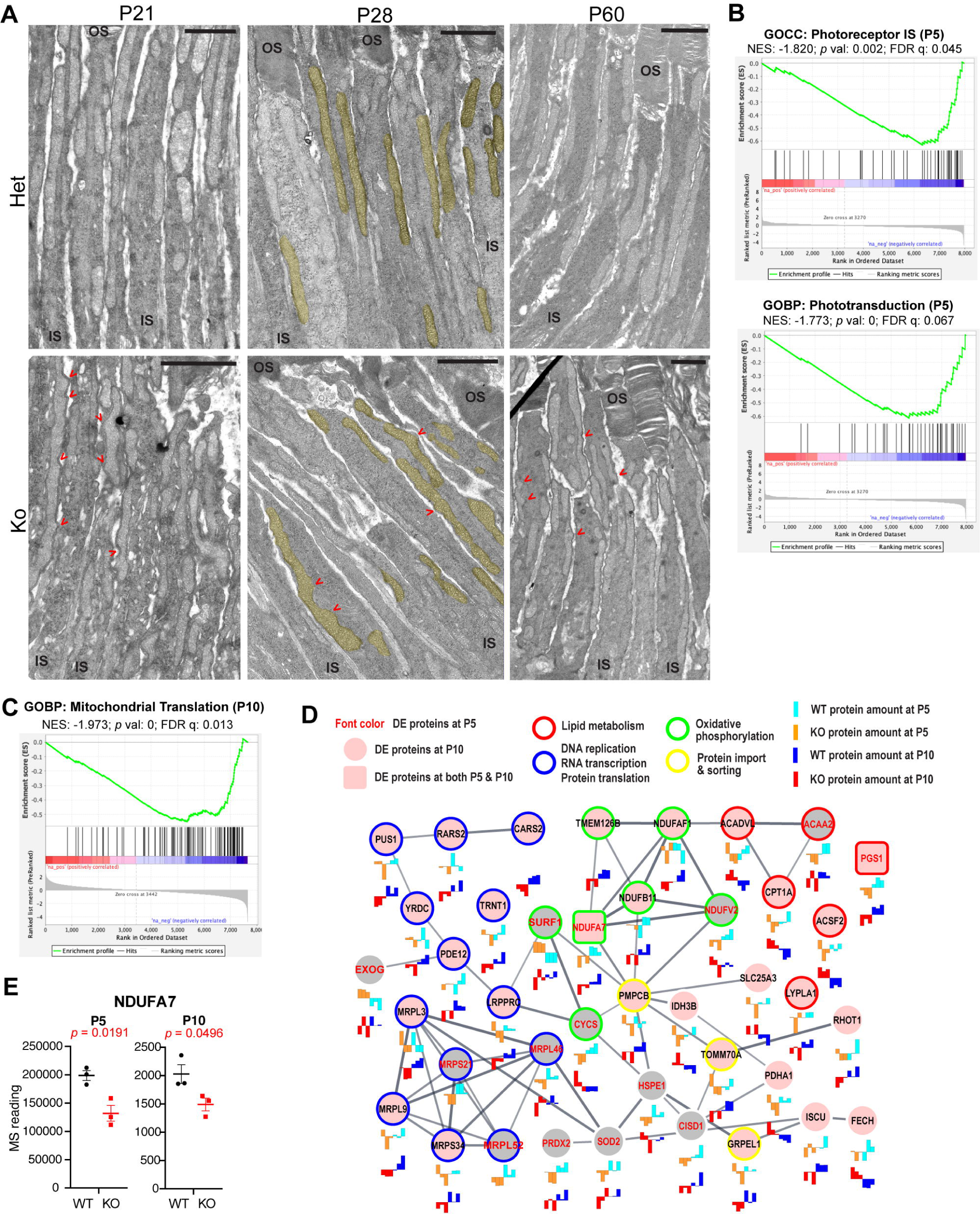
Mitochondrial morphology, protein translation, and function are affected in *Cfap418^-/-^* photoreceptors. **(A)** Longitudinal view of *Cfap418^-/-^* photoreceptors shows uneven diameters along the length of mitochondria at P21, P28, and P60, compared with the smooth straight long bar-shaped mitochondria in *Cfap418^+/-^* littermate photoreceptors. Mitochondria are highlighted in yellow in the P28 panels. Red arrows point to the abnormal constrictions and protruding bumps of the mitochondria. Scale bars, 1.5 μm. **(B)** Photoreceptor IS and phototransduction proteins are enriched in the down-regulated proteins in P5 *Cfap418^-/-^*retinas as revealed by GSEA. **(C)** GSEA shows mitochondrial translation proteins are reduced in P10 *Cfap418^-/-^* retinas. **(D)** The abundances of mitochondrial proteins in central dogma, oxidative phosphorylation, lipid metabolism, and protein import/sorting are either reduced or increased in *Cfap418^-/-^* retinas at P5 and P10. **(E)** The protein level of NDUFA7 is reduced in both P5 and P10 *Cfap418^-/-^* retinas. Data of individual mice, mean, and SEM are shown.

**Figure 5:**
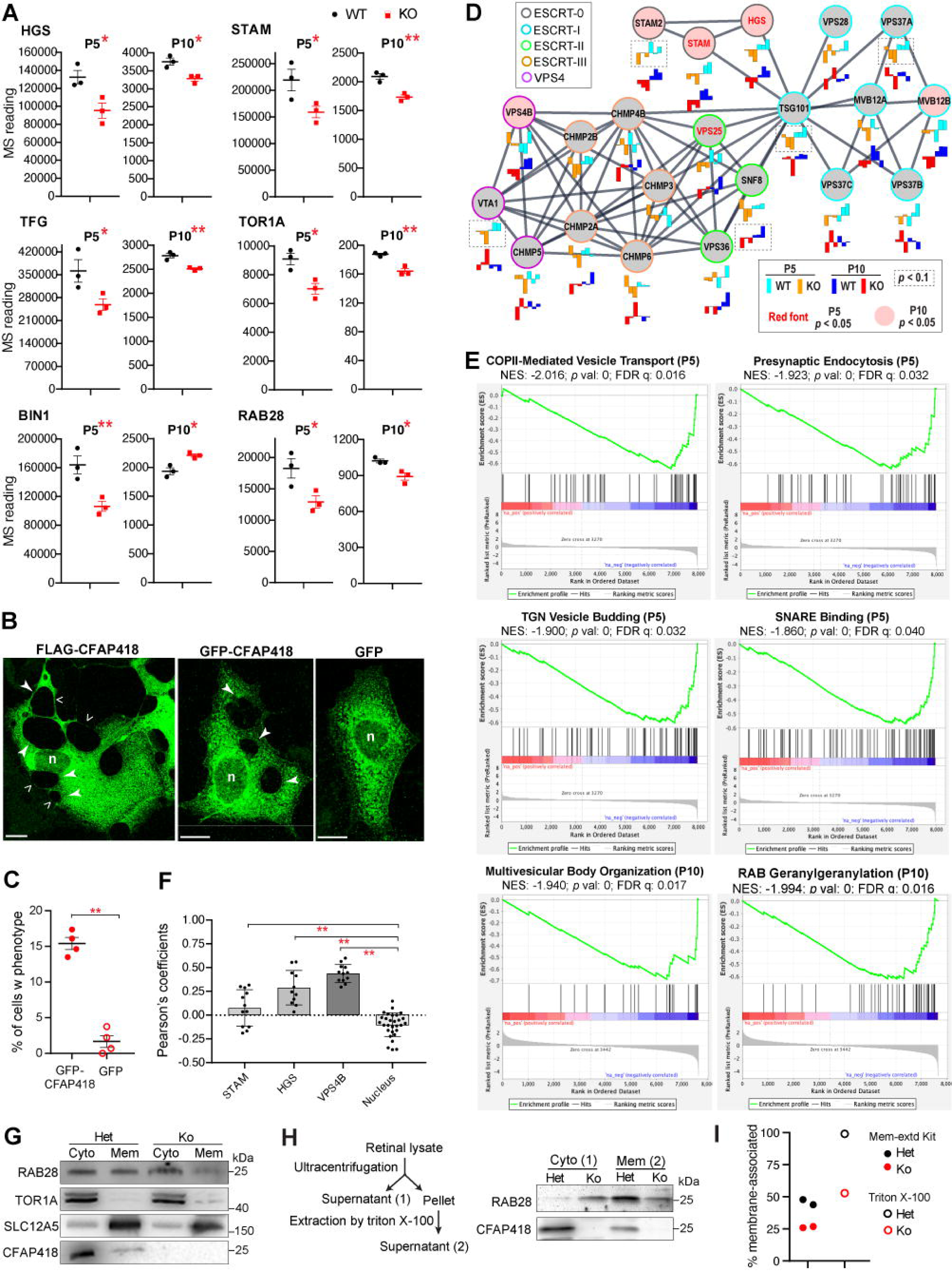
Membrane remodeling-associated proteins are altered at the onset of *Cfap418^-/-^*retinal phenotypes. **(A)** Six membrane remodeling-associated proteins are differentially expressed in P5 and P10 *Cfap418^-/-^*retinas. **(B)** Representative COS-7 cells transfected with FLAG- and GFP-CFAP418, but not GFP, accumulate large vacuoles (filled arrows). Open arrows: thin edges of vacuoles. n: nucleus. Scale bars: 10 μm. **(C)** Quantification of the vacuole accumulation phenotype in GFP-CFAP418- and GFP-transfected COS-7 cells. **(D)** Expressions of ESCRT complex components in *Cfap418^+/+^*and *Cfap418^-/-^* littermate retinas at P5 and P10. The expression levels of individual mice are shown as bar charts below each node. Lines between nodes: associations between nodes annotated by STRING /Cytoscape 3.8.1 (95, 96). **(E)** Representative vesicular trafficking processes that are negatively affected in P5 and P10 *Cfap418^-/-^* retinas, as revealed by GSEA. **(F)** PCCs of FLAG-CFAP418 with endogenous STAM, HGS, and VPS4B in COS-7 cells. **(G-H)** RAB28 is increased in the cytosol (Cyto) and decreased in the membrane (Mem) in *Cfap418^-/-^* retinas, compared with *Cfap418^+/-^* retinas at P21 (using a commercial membrane protein extraction kit, **G**) and P30 (using our triton X-100 protocol, **H**). TOR1A and SLC12A5 blots verified the separation between cytosol and membrane fractions. **(I)** Quantification of RAB28 membrane association. Dot plots show data from individual mice/experiments/cells, mean, and SEM (**A**, **C**, and **F**) or individual experiments (**I**). * and **, *p* < 0.05 and 0.01, respectively (Student’s t-test in **A** and **C**, Turkey’s multiple comparisons in **F**).

Gene Set Enrichment Analysis (GSEA) on P5 and P10 retinal proteomes was conducted to investigate whether mitochondrial function was abnormal in *Cfap418^-/-^* retinas. First, GSEA identified gene sets of photoreceptor IS proteins (*p* = 0.002; FDR q = 0.045) and phototransduction proteins (*p* = 0; FDR q = 0.067) were down-regulated at P5 and the gene set of ciliary membrane proteins including photoreceptor OS membrane proteins was down-regulated at P10 (*p* = 0; FDR q = 0.013) (Figures 4B, 7A, and Table S4), which were consistent with the phenotypes observed in mature *Cfap418^-/-^*retinas (16) and thus validated our GSEA results. GSEA further identified that proteins in mitochondrial translation were reduced in P10 *Cfap418^- /-^* retinas (Figure 4C and Table S4).

We then analyzed the expressions of all mitochondrial proteins in P5 and P10 *Cfap418^-/-^* retinal proteomes, according to a mammalian mitochondrial protein database MitoCarta3.0 (43). Sixty-six of the mitochondrial proteins were differentially expressed in *Cfap418^-/-^*retinas at either P5 or P10 (*p* < 0.05). Consistently, DE proteins functioning in mitochondrial central dogma (DNA replication, RNA transcription, and protein translation) were reduced in *Cfap418^-/-^* retinas (Figure 4D). Moreover, the DE proteins in oxidative phosphorylation and protein import and sorting pathways were mostly reduced. However, the DE proteins in lipid metabolism had mixed changes (Figure 4D). Besides PGS1 and ACSF2 mentioned above, CPT1A and ACADVL, responsible for the uptake of fatty acids and the first step of fatty acid beta-oxidation, were increased, while ACAA2, responsible for the last step of fatty acid beta-oxidation, was reduced, indicating that fatty acid beta-oxidation is disturbed. NADH:Ubiquinone oxidoreductase subunit 7 (NDUFA7), a subunit of complex I in the electron transport chain (44), was reduced in both P5 and P10 *Cfap418^-/-^* retinas (Figure 4E). Therefore, protein translation, oxidative phosphorylation, lipid metabolism, and protein import/sorting are abnormal in *Cfap418^-/-^* mitochondria. Based on the early onset of PGS1 and NDUFA7 changes at P5, the mitochondrial phenotype is likely one of the primary defects caused by membrane lipid disruption in *Cfap418^-/-^* photoreceptors.

### Membrane remodeling-associated proteins are affected in *Cfap418^-/-^* photoreceptor IS at ciliogenesis

Quantitative proteomic studies identified 11 DE proteins shared in both P5 and P10 *Cfap418^-/-^* retinas, besides PGS1 and NDUFA7 (Table S5). Six of them were associated with membrane remodeling in vesicular trafficking pathways. They were RAB28, HGS (hepatocyte growth factor-regulated tyrosine kinase substrate), STAM (signal transducing adaptor molecule), TFG (Trk-fused gene), BIN1 (bridging integrator-1), and TOR1A (torsin-1A) (Figure 5A). These proteins are involved in endosomal sorting complex required for transport (ESCRT)-mediated (RAB28, HGS, and STAM), BBSome-mediated ciliary (RAB28), ER to Golgi (TFG), ER to nuclear envelop (TOR1A), endosomal (BIN1) trafficking as well as ciliary extracellular vesicle/photoreceptor OS shedding (RAB28) (22, 45-53) (Table S5). These proteins were reduced in *Cfap418^-/-^* retinas at P5 and P10, except that BIN1 was reduced at P5 but increased at P10 (Figure 5A). The magnitude of these proteins’ changes was moderate in the range of 13% - 35%. However, the MS values of these DE proteins, except TOR1A at P10, were all above or around the median MS values in each animal. Thus, the MS values of these DE proteins were reliable. Additionally, our quantitative MS experiments were conducted on independent sets of retinas at P5 and P10 using two completely different protocols (label-free and TMT-labeling) by two proteomics core facilities. We considered that the DE proteins identified from both the P5 and P10 MS experiments were cross-confirmed. Because the DE proteins were identified in *Cfap418^-/-^*retinas as early as P5, when photoreceptor OS is not formed (54), the membrane-remodeling defects should occur in photoreceptor IS. Consistently, we observed that over-expression of FLAG- and GFP-CFAP418, but not GFP, in COS-7 cells resulted in large vacuole accumulation in the cytoplasm (Arrows, Figures 2B, 5B, 5C, and S3B), indicating a defect in membrane remodeling.

To further investigate the role of CFAP418 in membrane remodeling, we analyzed ESCRT proteins in the P5 and P10 retinal proteomes. Besides HGS and STAM, other ESCRT-0, ESCRT-I, ESCRT-II, and VPS4 proteins were also reduced (*p* < 0.05) or had a trend of reduction (*p* < 0.1) in P5 or P10 *Cfap418^-/-^* retinas (Figure 5D). Additionally, GSEA identified many down-regulated gene sets involved in vesicle budding, transport, targeting, and tethering, including the COPII-mediated vesicle transport and SNARE binding gene sets at P5 and multivesicular body (MVB) organization and RAB geranylgeranylation gene sets at P10 (Figure 5E and Table S4). These findings were supported by our previous observation of MVB accumulation in *Cfap418^-/-^*photoreceptor IS (16). We further found that CFAP418 was partially colocalized with VPS4B, HGS, and STAM in FLAG-CFAP418-transfected COS-7 cells (Figures 5F and S3A). However, VPS4B, HGS, and STAM did not interact with CFAP418 as shown in our AP-MS experiments (Table S1). Therefore, it is possible that CFAP418 binds to PA near these ESCRT proteins and disruption of this binding decreases ESCRT protein abundances.

To study whether the disruption of CFAP418 lipid binding affected membrane-protein associations, we focused on RAB28, which associates with membranes through prenylation (55). We separated RAB28 in retinal cytosol and membrane fractions using a commercial membrane protein extraction kit at P21 (Figure 5G) and our Triton X-100 extraction protocol at P30 (Figure 5H). Compared with *Cfap418*^+/-^ littermate retinas, RAB28 in membranes was reduced by ∼50% in *Cfap418*^-/-^ retinas (Figure 5I). Because only a small amount of RAB28 interacts with CFAP418 (Figure 1G) and is present, together with CFAP418, in photoreceptor IS (16, 22), the 50% reduction of RAB28 membrane association probably occurs in both IS and OS and results from the membrane lipid changes in *Cfap418*^-/-^ retinas, which is expected to affect the membrane biophysical and biochemical properties.

We further examined the distributions of ESCRT, endosomal, RAB28, and TFG proteins in *Cfap418^-/-^* photoreceptors by immunostaining. All these proteins were localized normally at P5. At P10, STAM and HGS were present as fine puncta in *Cfap418^+/-^* IS and OS, but they were enriched as large puncta in *Cfap418^-/-^* OS (Figures 6A and 6B). At P21, STAM and HGS were largely present in *Cfap418^+/-^* IS (Figures 6C and 6D), but they displayed reduced IS signals and prominent punctate OS signals in *Cfap418^-/-^* photoreceptors (Figures 6C and 6D). The distributions of STAM and HGS were however normal in *Cfap418^-/-^*photoreceptor OPL at P10 and P21. RAB28 immunoreactivity was diffusely distributed mainly in *Cfap418^+/-^* OS at P10 and P21 (Figures 6 and S4A). However, it became fragmented at the OS and RPE junction in *Cfap418^-/-^*photoreceptors at P21 (Figures 6E and 6F). The immunoreactive signal patterns of TFG, VPS4B, EEA1, RAB5, RAB7, and RAB11 were normal in *Cfap418^-/-^* photoreceptors at P10 and/or P21 (Figure S4). In summary, *Cfap418* knockout causes mislocalization of ESCRT-0 complex and RAB28 proteins in photoreceptors.

**Figure 6:**
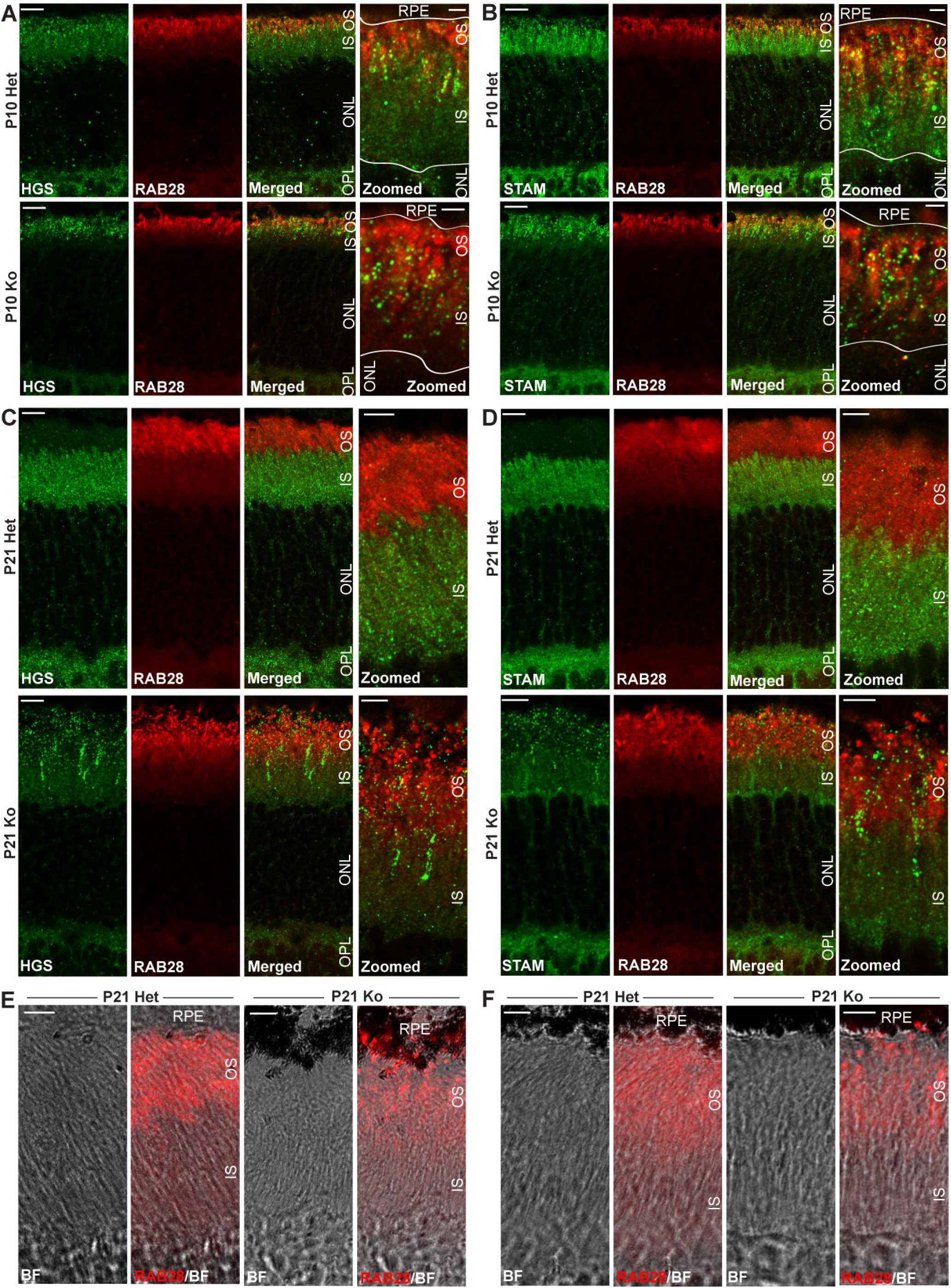
Abnormal HGS, STAM, and RAB28 distributions in *Cfap418^-/-^* photoreceptors. **(A)** HGS and **(B)** STAM, but not RAB28, signal patterns are different in P10 *Cfap418^+/-^* and *Cfap418^-/-^* photoreceptors. White lines outline the RPE and OS border and the IS and ONL border, based on the corresponding bright-field (BF) channels. **(C)** HGS and **(D)** STAM are mislocalized from the IS to the OS in P21 *Cfap418^-/-^* photoreceptors. **(C-F)** RAB28 signal shows an abnormal punctate pattern at the OS and RPE junction in P21 *Cfap418^-/-^* retinas. The zoomed images on the right in **A-D** are independent images from the ones on the left. The exact zoomed images in **C** and **D** are shown with RAB28 and BF channels in **E** and **F**, respectively. Scale bars: 2 μm (**A**, **B**, zoomed); 5 μm (**C**, **D**, zoomed; and **E**, **F)**; and 10 μm (others).

**Figure 7:**
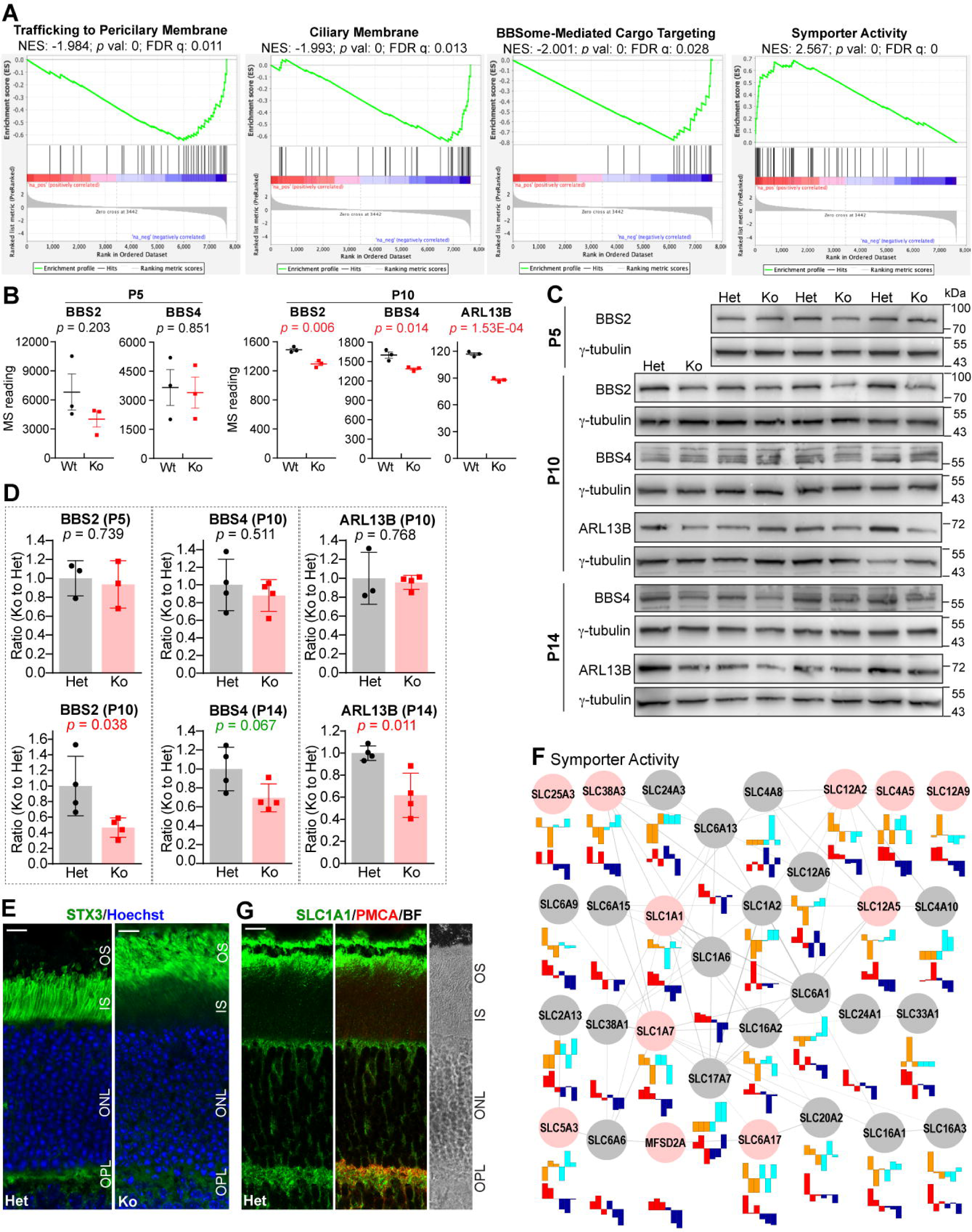
Ciliary transport proteins and symporters are affected during *Cfap418^-/-^*OS growth. **(A)** Proteins in several ciliary transport pathways are reduced, while symporters are increased in P10 *Cfap418^-/-^*retinas. **(B)** Quantitative MS data show normal BBS2, BBS4, and undetectable ARL13B protein expressions in P5 *Cfap418^-/-^* retinas but their reduced expressions in P10 *Cfap418^-/-^* retinas. **(C)** Semi-quantitative immunoblots for BBS2, BBS4, and ARL13B in *Cfap418^+/-^* and *Cfap418^-/-^* littermate retinas at different time points. The corresponding γ-tubulin immunoblots are loading controls. **(D)** Quantification of the semi-quantitative immunoblots reveals BBS2 and ARL13B reductions in P10 and P14 *Cfap418^-/-^* retinas, respectively, and a trend of BBS4 reduction in P14 *Cfap418^-/-^* retinas. **(E)** Mislocalization of STX3 from the IS and OPL to the OS in P21 *Cfap418^-/-^* photoreceptors. **(F)** The expressions of leading symporters, revealed by GSEA, in *Cfap418^+/+^* and *Cfap418^-/-^* retinas. Legends are the same as those in Figure 5D. **(G)** SLC1A1 is present in OS apex, ONL, and OPL in photoreceptors. OPL is marked by PMCA. Note that there is a tissue separation between OS and RPE in these images, as shown in the bright field (BF) channel. Scale bars, 10 μm. Data from individual mice and mean ± SEM are shown in **B** and **D**.

### Ciliary transport proteins and symporters are altered in *Cfap418^-/-^* photoreceptors after OS growth

GSEA identified several down-regulated pathways in P10 but not P5 *Cfap418^-/-^* retinas, which included the trafficking to the periciliary membrane, ciliary membrane components, and BBSome-mediated ciliary trafficking pathways (Figure 7A and Table S4). Among them, BBSome components (BBS proteins) and ARL13B were shared. Quantitative MS data showed that BBS proteins were normal at P5, but BBS2, BBS4, BBS5, BBS7, and ARL13B were reduced in a range of 13% to 25% in *Cfap418^-/-^* retinas at P10 (Figure 7B and Table S3). Because CFAP418 is associated with BBS disease (7), BBSome binds to PA (56, 57), and ARL13B associates with membranes through palmitoylation (58, 59), we performed semi-quantitative immunoblotting analysis for BBS2, BBS4, and ARL13B. BBS2 expression was normal at P5 but was reduced by ∼55% at P10 in *Cfap418^-/-^*retinas. The expressions of BBS4 and ARL13B were normal at P10 in *Cfap418^-/-^*retinas, but at P14, ARL13B was reduced by ∼40%, and BBS4 had a trend of reduction (Figures 7C and 7D). Therefore, the results from semi-quantitative immunoblotting analysis confirmed the findings from quantitative MS, although the former is less sensitive. Syntaxin 3 (STX3) is a cargo of the BBSome in photoreceptors (60-62). We found that STX3 was mislocalized from the IS and OPL to the OS in *Cfap418^-/-^* photoreceptors at P21 (Figure 7E). Together, we showed reductions of BBS and ARL13B proteins and dysfunction of the BBSome in *Cfap418^-/-^*retinas.

Symporter activity pathway was in a cluster of 45 overlapping up-regulated pathways in P10 *Cfap418^-/-^* retinas, revealed by GSEA and EnrichmentMap (63). This pathway had the highest normalized enrichment score (NES) and a close to zero FDR q value (Figure 7A and Table S4). Symporters are transmembrane proteins and co-transport two or more ions, amino acids, sugars, lipids, and neurotransmitters in the same direction across cell membranes (64). Forty-six symporters were detected in our P5 and P10 proteomes. Their expressions were normal in P5 *Cfap418^-/-^* retinas. However, 11 symporters were increased at P10 (Figure 7F). Among them, SLC1A7 and SLC12A2 were previously reported to localize in photoreceptor synaptic terminals (65, 66). Immunostaining for SLC1A1 showed that this symporter was localized at multiple photoreceptor layers at P10 and P21 (Figures S4B and 7G). SV2 and PMCA1 were used as OPL markers. There was no apparent difference in SLC1A1 distributions between *Cfap418^+/-^*and *Cfap418^-/-^* retinas at P10 and P21 (Figures S4A and not shown). Therefore, our results suggest that symporters’ expressions are likely increased in photoreceptors.

### Phosphorylation of proteins, including PA-binding protein kinase Cα, is altered in developing *Cfap418^-/-^* retinas

We analyzed protein phosphorylation in our P5 and P10 quantitative proteomic MS data (Tables S2 and S3). DPYSL3 (dihydropyrimidinase-related protein 3) was the only protein that was differentially phosphorylated (DP) in *Cfap418^-/-^*retinas at both P5 and P10 (Figure 8A). Compared with *Cfap418^+/+^* retinas, DPYSL3 phosphorylation was reduced by ∼35% at P5 but increased by ∼30% at P10, while the total DPYSL3 expression level was normal at both P5 and P10 (Figure 8B). The DP site of DPYSL3 at P10 was serine 101 (NP_001278384), which had not been well characterized.

**Figure 8:**
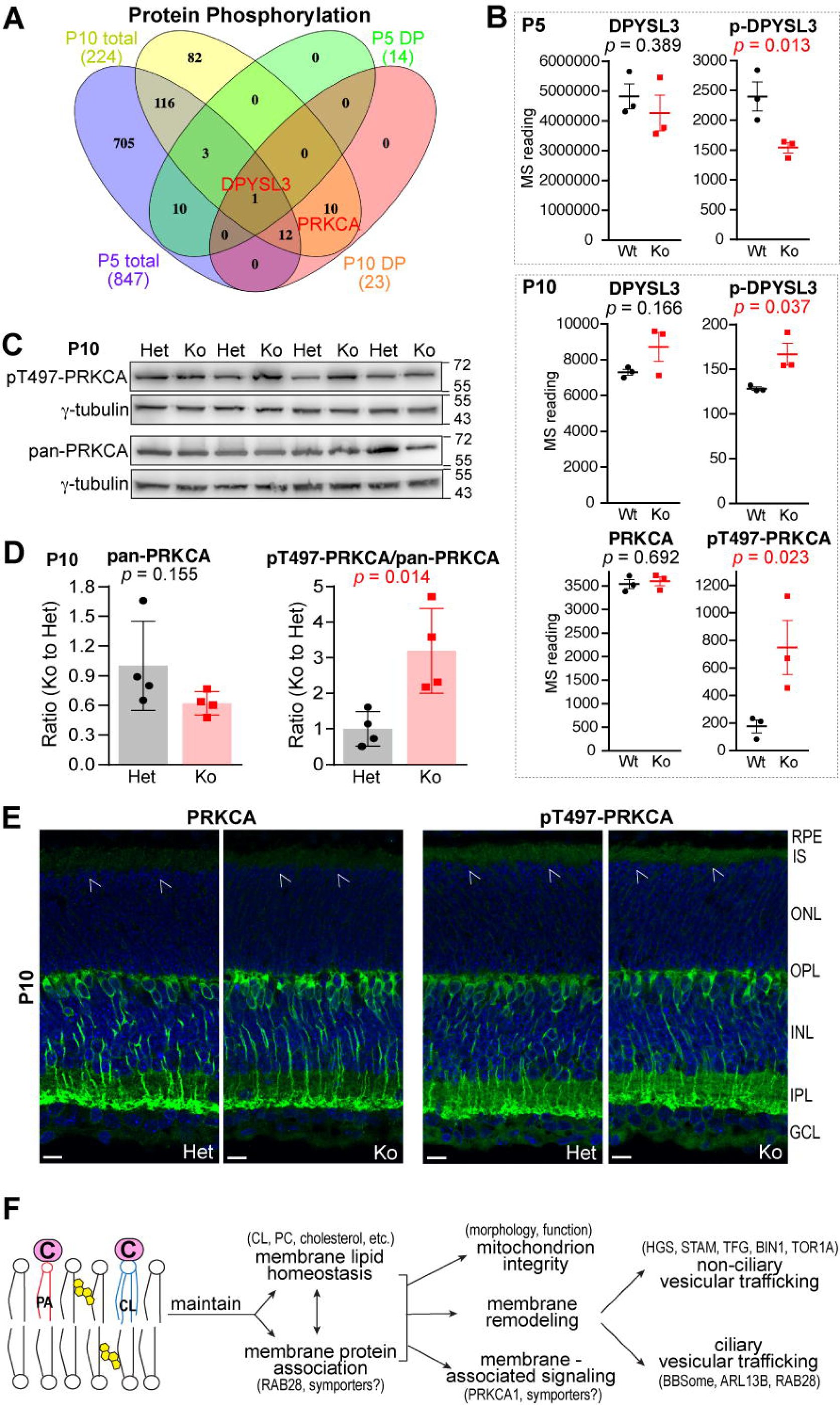
Protein phosphorylation is affected in developing *Cfap418*^-/-^ photoreceptors. **(A)** DPYSL3 is the only shared DP protein in P5 and P10 *Cfap418^-/-^* retinas, while PRKCA is a DP protein in P10 *Cfap418^-/-^* retinas. **(B)** Quantitative MS demonstrates that DPYSL3 phosphorylation is reduced in P5 *Cfap418^-/-^*retinas, and DPYSL3 and PRKCA phosphorylation is increased in P10 *Cfap418^-/-^*retinas. **(C)** Semi-quantitative immunoblots for pan- and pT497-PRKCA in 4 pairs of P10 *Cfap418^+/-^* and *Cfap418^-/-^* retinas. ψ-tubulin is a loading control. **(D)** Quantification of the semi-quantitative immunoblots for pan- and pT497-PRKCA. **(E)** Immunostaining displays similar pan- and pT497-PRKCA signal patterns between P10 *Cfap418^+/-^*and *Cfap418^-/-^* littermate retinas. The pT497-PRKCA signal is stronger in photoreceptors than the pan-PRKCA signal (arrows). Scale bars, 10 μm. **(F)** Proposed model for the CFAP418 molecular function in cells. Data from individual mice and mean ± SEM are shown in **B** and **D**.

At P10, the phosphorylation of protein kinase Cα at threonine 497 (pT497-PRKCA) was increased 4.3-fold in *Cfap418^-/-^* retinas, while the total PRKCA expression was unchanged, compared with *Cfap418^+/+^* retinas (Figure 8B). This phosphorylation site is critical for PRKCA activation (67) and was the only DP site that had an antibody commercially available among all the identified DP proteins/sites at P5 and P10. Semi-quantitative immunoblot analysis was conducted. After normalization by loading control γ-tubulin, the ratio of pT497-PRKCA to pan-PRKCA signal was increased 3.2-fold in *Cfap418^-/-^*retinas, compared with *Cfap418^+/-^* retinas (Figures 8C and 8D). Because PRKCA was localized only to the rod bipolar cells in mature rodent retinas (Figure S5) (68, 69), we examined PRKCA distribution in P5 and P10 mouse retinas by immunostaining. Both PRKCA and pT497-PRKCA were localized throughout different retinal layers at P5 (Figure S5). At P10, PRKCA and especially pT497-PRKCA were present in photoreceptor IS and rod bipolar cells (Figure 8E). Their signal patterns were the same between *Cfap418^+/-^* and *Cfap418^-/-^* retinas (Figure 8E). Therefore, as a PA-interacting protein (70), PRKCA’s activity, indicated by its phosphorylation at T497, is enhanced probably in both *Cfap418^-/-^* photoreceptors and rod bipolar cells during development.

## Discussion

We investigated the CFAP418 molecular function in the retina using several unbiased omics approaches combined with traditional biological methodology. We unexpectedly discovered that CFAP418 binds directly to PA and CL on cell membranes but does not bind strongly to specific proteins. The disruption of CFAP418 bindings to lipids leads to extensive changes in membrane lipid composition and weakens RAB28 membrane association. Upon ciliogenesis, *Cfap418* knockout reduces CL synthase PGS1 and membrane remodeling-associated proteins, which is followed by compromised mitochondrial morphology and function and mislocalization of ESCRT-0 proteins HGS and STAM in photoreceptors. *Cfap418* knockout also reduces ciliary transport BBSome and ARL13B proteins; disturbs BBSome function; increases transmembrane symporters and PRKCA activity during photoreceptor OS growth (Figure 8F). In summary, we have identified CFAP418 as a novel lipid-binding protein, which is critical for membrane lipid homeostasis and a series of membrane-associated cellular events. Although CFAP418 is named a ciliary and flagellar associated protein, this protein functions in non-ciliary compartments, which contributes to ciliogenesis indirectly. This study sheds new light on the multiple roles of membrane lipid homeostasis in cells, which has been severely understudied, and demonstrates that disruption of the membrane lipid homeostasis is a novel pathological mechanism underlying inherited retinal degenerations and syndromic ciliopathies. We further show that the application of combined various omics approaches is an effective and efficient way to reveal the molecular function of little characterized disease genes, especially the genes that are related to membrane lipid homeostasis and challenging to study. Our results open a new avenue to further mechanistic studies on membrane lipid biology, ciliogenesis, ciliopathies, and other lipid-related genetic diseases.

The direct interactions of CFAP418 with PA and CL but no strong and static interaction of CFAP418 with proteins indicate that the membrane lipid imbalance occurs upstream of the protein expression and phosphorylation changes in *Cfap418*^-/-^ retinas, although our lipidomic study was conducted at P10 later than our label-free proteomic study at P5. PA is a precursor of many glycerophospholipids in membranes and glycerolipids in lipid droplets (26) and is a hub for membrane biogenesis and lipid storage (71). The decreases in many glycerophospholipids and increases in some DAG and TAG in *Cfap418*^-/-^ retinas (Fig. 3C) suggest an imbalance between membrane biogenesis and lipid storage. The widespread changes across different lipid categories in *Cfap418*^-/-^ retinas are probably due to the interconnection of the metabolic pathways of lipid categories. These changes are expected to affect the membrane biophysical and biochemical properties and the abundance and function of many integral and peripheral membrane proteins. This notion is supported by our observations of decreased RAB28 membrane association and altered HGS, symporter, and photoreceptor OS membrane protein expressions in *Cfap418*^-/-^ retinas. PA is cone-shaped with a small head group. It induces negative membrane curvature and actively participates in membrane fission and fusion (3). Consistently, we observed membrane remodeling phenotypes in both CFAP418 overexpressed cells and CFAP418 knockout retinas. In line with the knowledge that PA interacts with proteins to stabilize protein conformation and regulate protein function (3, 72), we discovered an activity decrease and increase of PA-binding BBSome and PRKCA, respectively, in *Cfap418*^-/-^ retinas. Finally, CL is present in mitochondrial membranes only, where it is synthesized from and hydrolyzed to PA (26). Both PA and CL play a role in mitochondrial dynamics and function (23, 73). Thus, we observed mitochondrial morphological and functional defects in *Cfap418*^-/-^ photoreceptors.

However, many questions remain unanswered. For example, it is unclear whether CFAP418 functions as a sequester to store PA and CL in specific membrane domains, a regulator for PA and CL synthesis and their production of other lipids, a modulator of PA and CL roles in protein activities, and/or something else. It is undetermined whether CFAP418 loss changes PA abundances and composition in cell membranes. Additionally, there are no consensus PA- and CL-binding motifs in proteins (3, 72, 74, 75). Hydrophobic and positively charged residues have been proposed to participate in PA- and CL-binding. The PA-binding of these residues is proposed mediated via a hydrogen bond switch. CFAP418 is enriched with positively charged residues. But the mechanisms and modes of CFAP418 bindings to PA and CL are unknown. The bindings of CFAP418 to PA and CL could be independent or rely on each other positively or negatively.

In mature *Cfap418*^-/-^ retinas, the photoreceptor OS exhibits the most evident phenotypes (16). Here, we discovered several defects during OS development that contribute to the phenotypes. First, the BBSome and ARL13B transport transmembrane and lipidated proteins into and out of the OS (58, 59, 61, 76, 77). We found reductions of BBS and ARL13B proteins and dysfunction of BBSome in developing *Cfap418*^-/-^ photoreceptors. Second, BBSome cargos RAB28 and STX3 function in cone OS shedding and OS membrane protein trafficking, respectively (22, 45, 46, 60, 78). We observed RAB28 expression and membrane association reduction and STX3 mislocalization in developing *Cfap418*^-/-^ photoreceptors. Third, the endosomal pathway is implicated in photoreceptor OS protein transport (79). In particular, HGS participates in cone PRPH2 OS trafficking (79). In *Cfap418*^-/-^ photoreceptors, both ESCRT-0 proteins HGS and STAM are mislocalized. Fourth, removal of docosahexaenoic acid (DHA, 22:6) in glycerophospholipids disrupts photoreceptor OS disc alignment (80), a phenotype similar to that in mature *Cfap418*^-/-^ photoreceptors (16). DHA also affects STX3 conformation and function in photoreceptors (81). We discovered reductions of many glycerophospholipids containing the DHA acyl chain in developing *Cfap418*^-/-^ retinas. Fifth, the abnormal ratio of PC to PE and ratio of longer polyunsaturated to shorter saturated acyl chains in *Cfap418*^-/-^ retinas could result in protein mislocalization across the center and rim of OS membrane discs (82). Although PRKCA participates in endocytic membrane trafficking and can be regulated by membrane lipids other than DAG (83, 84), the function of PRKCA and most DE symporter proteins in developing photoreceptors have not been studied and the physiological relevance of their changes in *Cfap418*^-/-^ photoreceptors remains to be elucidated.

Some lipid metabolic defects may explain the protein expression and activity changes in *Cfap418*^-/-^ photoreceptors. For example, HGS binds to PI(3)P via its FYVE domain (49). In *Cfap418*^-/-^ retinas, the reduction of PIK3CA, which phosphorylates PI to generate PI(3)P, and the increase of MTMR2, which dephosphorylates PI(3)P to generate PI, are expected to decrease PI(3)P perhaps in photoreceptor IS, which may lead to the HGS mislocalization to photoreceptor OS, where the discs are enriched with PI(3)P (85). Furthermore, the increase of PRKCA activity may be related to the increases of PLCD1 expression and DAG abundance in *Cfap418*^-/-^ retinas. While these connections between lipid and protein defects need to be further verified in *Cfap418*^-/-^ photoreceptors, it would be interesting to understand how the affected lipids and proteins coordinate and function together to ensure the proper vesicular trafficking, especially the ciliary trafficking of OS integral and peripheral membrane proteins, in healthy photoreceptors.

The combined application of quantitative lipidomic, proteomic, phosphoproteomic, and bioinformatic approaches (63, 86-89) enabled us to systematically integrate the *Cfap418*^-/-^ phenotypes from different angles. Using a large sample size in our untargeted lipidomic study significantly increased the sensitivity to detect changes in lipid species. These changes can be explained by the expression alterations in some membrane lipid metabolic enzymes and transporters revealed by our proteomic data, although the activities of these proteins were unable to assess. Interestingly, the increase in cholesterol and decrease in PC with DHA acyl chains discovered in our *Cfap418*^-/-^ retinas were also found in the photoreceptor OS of 5-month-old *bbs1*^-/-^ zebrafish (90), a BBS model at a later disease progression time point. In addition to the reliability, objectivity, and broad coverage, our quantitative proteomics approaches are more sensitive than the traditional semi-quantitative immunoblot analyses. In general, moderate expression and phosphorylation changes were seen in our DE and DP proteins. These changes complement each other. Some of them were confirmed by independent techniques or highly consistent with known *Cfap418*^-/-^ phenotypes. Besides the 8 DE proteins mentioned in this report, 5 more DE proteins are shared in P5 and P10 proteomes (Table S5). The membrane association of these proteins and the physiological significance of their changes are unclear. Additionally, hundreds of other DE proteins, tens of DP proteins, and tens of altered molecular functions and cellular pathways were identified in this study (Tables S3 and S4). A detailed investigation of these alterations will generate a more complete picture regarding the role of CFAP418 and membrane lipid homeostasis at various membrane structures and vesicular transport pathways. Among the known proteins associated with IRD diseases (RetNet), 25 are DE proteins in P5 and/or P10 *Cfap418*^-/-^ retinas (Figure S6). Therefore, our omics data are valuable resources for studying proteins, cellular pathways, and human diseases associated with CFAP418 and its related proteins.

In summary, our findings demonstrate that CFAP418, as a lipid-binding protein, maintains lipid homeostasis in many cellular membranes and that the integrity of cellular membranes is essential for mitochondrial morphology and function, non-ciliary and ciliary vesicular trafficking pathways, and membrane-associated signaling. Our studies reveal the pathogenic mechanism underlying the defects in photoreceptors and perhaps other ciliated cells caused by *CFAP418* mutations and indicate that this mechanism is partially shared with other IRD genes, e.g., the BBS and RAB28 genes. Considering its high evolutionary conservation, expression in many developing and mature tissues (7, 14), and involvement in melanosome transport in non-ciliated zebrafish melanophores (7), the role of CFAP418 in lipid binding, homeostasis, and membrane remodeling is likely conserved among species and tissues and is important for cell survival and function.

## Methods

### Mice and cell lines

*Cfap418*^-/-^ mice with a mixed CBA/C57BL6 genetic background have been described previously (16). These mice were maintained, cared for, and examined according to the protocol approved by the Institutional Animal Care and Use Committee at the University of Utah. Mice of both sexes were randomly assigned to experimental groups at P5, P10, P21, and P30. Littermate *Cfap418*^+/+^ or *Cfap418*^+/-^ mice were included as controls. HEK293 (CRL 10852; RRID:CVCL_6974) and COS-7 (CRL 1651; RRID:CVCL_0224) cell lines were purchased from ATCC (https://www.atcc.org). These cells were cultured in Dulbecco’s modified Eagle’s medium supplemented with 10% (v/v) fetal bovine serum, 50 unit/ml penicillin, and 50 μg/ml streptomycin (ThermoFisher Scientific, Waltham, MA). One Shot™BL21 Star™ (DE3) cell line was purchased from ThermoFisher Scientific (C601003) and cultured according to the manufacturer’s protocol.

### Pulse-chase experiment

Pulse-chase experiments were conducted at 37°C in a humidified 5% CO2 incubator. Four sets of three retinas per genotype were incubated in DMEM without methionine or cysteine (ThermoFisher Scientific. cat# 21013) but supplemented with 200 mM cysteine (SigmaAldrich C7352) for 30 minutes. The medium was then changed to DMEM with cysteine and 0.05 μCi/μl ^35^S methionine (PerkinElmer, cat. NEG009A). Pulse labeling was done for 30, 60, 90, or 120 minutes. For the chase experiment, the radioactive medium was replaced with cold DMEM, and incubation was carried out for 3 hours or overnight. The radiolabeled retinas were stored at -80°C. The frozen retinas were subsequently homogenized by repetitive pipetting on ice in PBS with 0.5% TritonX-100 and cOmplete protease inhibitors (Roche). The homogenates were precleared by incubation with protein G Sepharose (GE Healthcare, cat#17-0618-02) for 1 hour at 4°C. After centrifugation, the supernatants were incubated overnight at 4°C with Protein G-bound antibodies, which were generated by mixing antibodies with Protein G in PBS and 1% TritonX-100 for 2 hours at 4°C. Following the incubation, samples were washed three times with wash buffer (50 mM Tris pH7.5, 0.3 M NaCl, 0.1% TritonX-100, 5 mM EDTA) and once with PBS. The final pellets were extracted by adding protein gel loading buffer, run on a 10% SDS-polyacrylamide gel, dried, exposed to a storage phosphor screen, and scanned by an S35 Typhoon Trio Phosphoimager (GE Healthcare).

### RT-qPCR

RT-qPCR was conducted as previously described (16). Briefly, retinal total RNA was isolated using the SurePrep RNA purification kit (Fisher Scientific, Hampton, NH) and reverse transcribed using the ThermoScript RT-PCR system (ThermoFisher Scientific, Waltham, MA). qPCR was performed using the SYBR Premix Ex Taq kit (TaKaRa Bio USA, San Jose, CA) and the CFX Connect real-time PCR machine (Bio-Rad, Hercules, CA). The primer information is shown in the Key Resources Table. The PCR condition was 95°C for 2 min followed by 45 cycles of 95°C for 10 sec and 55°C for 30 sec (60°C for IRE1α). *C*q was determined using the single threshold mode of the CFX Manager software.

### Lipid binding experiment

His- and GST-tagged CFAP418 proteins were expressed in One Shot™ BL21 Star™ (DE3) cells (ThermoFisher, Waltham, MA) and purified using HisPur™ Ni-NTA resin and Pierce™ Glutathione Superflow Agarose, respectively (ThermoFisher, Waltham, MA). After spotting 1 ng of the purified recombinant proteins on the membrane lipid and PIP strips (P-6002 and P-6001, respectively, Echelon Biosciences, Salt Lake City, UT) as a positive control, the strips were blocked in 3% bovine serum albumin in PBS-T buffer for 1 hour and then incubated in the same buffer with the recombinant proteins at a concentration of 1 μg/ml. The bound CFAP418 protein was detected by sequential incubations with CFAP418 antibody (16) and a horseradish peroxidase-conjugated secondary antibody. The enhanced chemiluminescence was scanned by a FluorChem Q machine (Cell Biosciences, Inc., Santa Clara, CA) or an iBright™ CL750 imaging system (ThermoFisher, Waltham, MA).

### Transmission electron microscopy

Mouse eyeballs were enucleated and briefly fixed in 1% formaldehyde and 2.5% glutaraldehyde in 0.1 M cacodylate buffer, pH 7.5. After removal of the cornea and lens, the obtained eye cups continued to be fixed overnight. The post-fixation with 2% osmium tetroxide, dehydration with a graded alcohol series, embedding in Epon, sectioning with ultramicrotome, and post-staining with lead citrate were performed as described previously (16). The transmission electron micrographs were taken using a JEOL JEM-1400 TEM equipped with a Gatan 4000 camera or a Gatan Orius SC1000B camera.

### Immunoblotting, immunoprecipitation, and immunostaining

Immunoblotting, immunoprecipitation, and immunostaining procedures were described previously with minor changes (91). Briefly, mouse retinas were homogenized by pipetting up and down using a wide-opening pipet tip and lysed at 4°C for 30 minutes in a solubilization buffer (50 mM Tris-HCl, pH 7.5, 100 mM NaCl, 5 mM EDTA, 1% Triton X-100, 0.05% SDS, 2.5% glycerol, and 1.0 mM phenylmethylsulphonyl fluoride). The retinal lysates were centrifuged at 14,000 x g for 12 minutes. The SDS sample loading buffer was added to the supernatants, which were run on a 10% or 12% SDS-PAGE. To separate retinal cytosol and membrane fractions, the retinal lysate, cleared by centrifugation at 14,000 x g for 10 minutes, was ultracentrifuged at 100,000 x g for 30 minutes. The membrane proteins were extracted from the pellet using lysis buffer containing 1% Triton X-100. Alternatively, the retinal cytosol and membrane proteins were separated using the Mem-PER™Plus membrane protein extraction kit (Cat #: 89842, ThermoFisher Scientific, Waltham, MA). The SDS sample loading buffer was added to the cytosol and membrane fractions, which were run on a 10% SDS-PAGE. The immunoblotting signals were developed using ProSignal® Pico ECL Reagent (Genesee Scientific, San Diego, CA), scanned by a FluorChem Q machine (Cell Biosciences, Inc., Santa Clara, CA) or an Invitrogen™ iBright™ CL750 machine (ThermoFisher Scientific, Waltham, MA), and quantified using ImageJ. The intensity of the immunoblotting signals was normalized by loading control γ-tubulin signals in the same lanes. The immunoprecipitation procedure was described in the affinity purification section. For cultured cells, immunoprecipitation was performed at 24 hours post-transfection.

For immunostaining, mouse eyeballs were enucleated and fixed briefly in 4% formaldehyde in phosphate-buffered saline (PBS). After removal of the cornea and lens, the eyecups were continued to be fixed for 1 hour before cryosection at 12 μm using Leica CM3050S cryostat. COS-7 and HEK293 cells were transfected with DNA plasmids using PEI (Polysciences, Inc., Warrington, PA). At 24 hours post-transfection, cells were fixed by methanol:acetone (19:1) at - 20°C for 10 min. The fixed retinal sections and transfected cells were then subjected to immunostaining procedures. The immunofluorescence signals were captured using a Leica SP8 confocal microscope with a HC PL APO 63X1.40 OIL CS2 objective. The colocalization of proteins in cultured cells was analyzed using the Coloc module of Imaris 9.7.2 and 9.8.2 (Oxford Instruments, Concord, MA). To quantify the vacuole accumulation phenotype, transfections of COS-7 cells with GFP-CFAP418 and GFP were conducted four times. Cells with vacuole accumulation were counted from 135 to 360 transfected cells in each experiment by a person blind to treatment. The dilution ratios of primary and secondary antibodies for immunoblotting, immunoprecipitation, and immunostaining were based on manufacturers’ suggestions.

### Quantitative proteomic, phosphoproteomic, lipidomic analyses, and affinity purification-mass Spectrometry

Details of various quantitative MS and AP-MS were described in supplemental methods. Proteins detected by label-free or TMT-labeling quantitative MS were ranked based on their values of -log(*p* value)*sign(log(FC)). For each protein, the *p* value was calculated using the normalized MS values generated by MetaboAnalyst 5.0 (92). The normalization parameters in MetaboAnalyst were set as normalization to constant sum; log10 transformation, and mean centering. The ranked proteins were then analyzed using the Gene Set Enrichment Analysis software (GSEA desktop v4.1.0) and the gene set databases of c2.cp.reactome.v7.4.symbols.gmt and c5.go.v7.4.symbols.gmt. The Run GSEAPreranked tool with default basic and advanced parameters were applied except that the gene set size was set to 15-200 and the mouse gene symbols were remapped to human orthologs. At P5, 21 and 1 out of the 4,033 screened gene sets were down-regulated and up-regulated in *Cfap418^-/-^* retinas, respectively (FDR q < 0.05, Table S4). At P10, 24 and 86 out of the 3988 screened gene sets were down-regulated and up-regulated in *Cfap418^-/-^* retinas, respectively (FDR q < 0.05, Table S4). The up- and down-regulated gene sets were visualized and further analyzed by EnrichmentMap and AutoAnnotate using Cytoscape 3.8.1 (63). Only one entry of proteins identified multiple times in each proteomic experiment was considered in the Venn Diagram analyses. Venn Diagrams for the quantitative proteomic and phosphoproteomic MS data were drawn using VENNY 2.1 (https://bioinfogp.cnb.csic.es/tools/venny/). The Venn Diagrams for the affinity purification/MS data were drawn using InteractiVenn (93).

### Statistics

Protein and phosphoprotein intensities generated from label-free and TMT-labeling MS were processed using the Statistical Analysis module in MetaboAnalyst 5.0 (89, 92). The wild-type and *Cfap418^-/-^* samples were normalized by their own sum, converted to an approximately normal distribution by logarithmic transformation (base 10), and scaled by mean centering. Principal component analysis (PCA) and heatmap hierarchical clustering with default parameters were implemented to examine the data quality and detect the potential outliers. The t-test in the Univariate Analysis category was conducted to identify differentially expressed (DE) and phosphorylated (DP) proteins. Because very few proteins had an FDR value (adjusted *p* value for multiple comparisons) smaller than 0.05, we defined DE and DP proteins as long as their raw *p* values were smaller than 0.05. There was no fold change cutoff for DE proteins at P5 or DP proteins at P5 or P10, while the fold change cutoff was 1.1 for the DE proteins at P10. The fold change was calculated by the ratio of *Cfap418^-/-^* protein/phosphoprotein intensity to wild-type protein/phosphoprotein intensity.

Lipidomic data were also processed and analyzed in MetaboAnalyst 5.0 (89, 92). The *Cfap418^+/-^* and *Cfap418^-/-^* samples were normalized by their own sum and scaled by mean centering without any data transformation. PCA and heatmap hierarchical clustering detected sample 14 (*Cfap418^-/-^*) as an outlier (Figure S2B and not shown). This sample was excluded from downstream analyses. Differentially abundant lipid species were identified by t-test with an FDR value smaller than 0.05 and a ratio of *Cfap418^-/-^* to *Cfap418^+/-^* value larger than 1.05 or smaller than 0.95. Lipid enrichment analysis was implemented using the Quantitative Enrichment Analysis (QEA) algorithm offered by the Metabolite Set Enrichment Analysis (MSEA) module. The metabolite set library of 1072 sub-chemical class metabolite sets was selected.

The results of RT-qPCR and semi-quantitative immunoblotting analyses as well as the abundances of different lipid categories and acyl chains were compared between *Cfap418^-/-^* and control (*Cfap418^+/+^* or *Cfap418^+/-^*) groups using Student’s t-test in Microsoft Excel.

## Author Contributions

AMC and DY designed and performed the experiments. DZ, GN, JAM, TB performed the experiments. JY designed the experiments, analyzed the data, and wrote the manuscript.

## Acknowledgments

The authors thank the Mass Spectrometry Proteomics Core at Baylor College of Medicine, Thermo Fisher Center for Multiplexed Proteomics and Taplin Mass Spectrometry Facility at Harvard Medical School, and Metabolomics Core at the University of Utah for conducting mass spectrometry experiments. The authors are grateful to the Bryan Jones laboratory, Ning Tian, Junhuang Zou, Guoxin Ying, and Ali S. Sharif at the University of Utah for their assistance with TEM, Imaris usage, confocal imaging, RAB28 constructs, and RT-qPCR, respectively. This work was supported by National Institutes of Health grants EY026521 (J.Y.), EY014800 (core grant to the Department of Ophthalmology, University of Utah), P30CA125123 (Baylor College Proteomic Facilities), International Retinal Research Foundation (J.Y.), Research to Prevent Blindness, Inc. (the Department of Ophthalmology at the University of Utah), and Cancer Prevention and Research Institute of Texas (RP170005, the Mass Spectrometry Proteomics Core at Baylor College of Medicine). The funders had no role in study design, data collection and analysis, decision to publish, or manuscript preparation.

## Notes

### Competing Interest Statement

The authors have declared no competing interest.

