## Supplemental Material for "Disruption of CFAP418 interaction with lipids causes abnormal membrane-associated cellular processes in retinal degenerations"

**KEY RESOURCES TABLE**

| **REAGENT OR RESOURCE** | **SOURCE** | **IDENTIFIER** |
|---|---|---|
| **Antibodies** | | |
| Rhodopsin (1D4) | Robert Molday, University of British Columbia | |
| DPYSL2 | This paper | |
| GNAT1 | Santa Cruz | sc-389; RRID:AB_2294749 |
| ATF6 | NovusBiologicals | 70B1413.1 |
| Phospho-IRE1α | NovusBiologicals | NB100-2323; RRID:AB_10145203 |
| γ-tubulin | Sigma-Aldrich | T6557; RRID:AB_477584 |
| RAB28 | Biorbyt | Orb11632 |
| CFAP418 | Custom-made (1) | |
| FLAG | Sigma-Aldrich | F1804; RRID:AB_262044 |
| KDEL | Abcam | ab12223; RRID:AB_11127118 |
| Golgi58K | Abcam | ab6284; RRID:AB2041471 |
| RAB5A | CellSignalingTech | 46449; RRID:AB_2799303 |
| RAB7 | CellSignalingTech | 9367; RRID:AB_1904103 |
| RAB11 | CellSignalingTech | 5589; RRID:AB_10693925 |

| | | |
|---|---|---|
| EEA1 | ThermoFisher | MA5-14794;<br>RRID:AB_10985824 |
| NDUFA7 | NovusBiologicals | NBP3-03240 |
| STAM | Proteintech | 12434-1-AP;<br>RRID:AB_2199965 |
| HGS | Proteintech | PA5-27491;<br>RRID:AB_2544967 |
| VPS4B | Proteintech | 17673-1-AP;<br>RRID:AB_2215504 |
| BBS2 | Santa Cruz | Sc-365355;<br>RRID:AB_10846953 |
| BBS4 | ThermFisher | PA3-912;<br>RRID:AB_10596774 |
| ARL13B | Proteintech | 177711-1-AP;<br>RRID:AB_2883085 |
| STX3 | Proteintech | 15556-1-AP;<br>RRID:AB_2198667 |
| SLC1A1/EAAC1 | ThermoFisher | 12686-1-AP;<br>RRID:AB_2188001 |
| PRKCA | AVIVA | OAAN00067 |
| Phospho-PRKCA | ThermoFisher | PA5-78125;<br>RRID:AB_2736409 |

| | | |
|---|---|---|
| TOR1A | ThermoFisher | PA5-65651;<br>RRID:AB_2662036 |
| PMCA | EMDMillipore | MABN1802 |
| mCherry | Rockland | 200-301-379;<br>RRID:AB_2611063 |
| TFG | ThermoFisher | 11571-1-AP;<br>RRID:AB_2203102 |
| SV2 | DSHB | SV2; RRID:AB-2315387 |
| Goat Anti-Mouse HRP | Thermofisher | 31432; RRID:AB_228302 |
| Goat Anti-Rabbit HRP | Thermofisher | G-21234; RRID:AB_1500696 |
| Donkey Anti-Goat HRP | Thermofisher | PA-28664;<br>RRID:AB_10990162 |
| AlexaFluor A488<br>Goat Anti-Rabbit | Thermofisher | A11034; RRID:AB_2576217 |
| AlexaFluor A488<br>Goat Anti-Mouse | Thermofisher | A11029; RRID:AB_2534088 |
| AlexaFluor A568<br>Goat Anti-Rabbit | Thermofisher | A11036; RRID:AB_10563566 |
| AlexaFluor A568<br>Donkey Anti-Goat | Thermofisher | A11057;RRID:AB_142581 |
| Hoechst 33342 | BD Biosciences | 561908; RRID:AB_2869394 |
| **Oligonucleotides** | | |
| PERK-forward: | This paper | N/A |

| | | |
|---|---|---|
| CAC AGG GAC CTC AAG CCT TCC | | |
| PERK-reverse: GGA CTC GTT CCA TCT GGG TGC | This paper | N/A |
| IRE1α-forward: GAC GAC GTGGACTACAAGATG | This paper | N/A |
| IRE1α-reverse: GGC GTT AGC TTG CTC TTG GCC | This paper | N/A |
| ATF6-forward: GGG TTC GGA TAT CGC TGT GCT G | This paper | N/A |
| ATF6-reverse: GCT CAT GGG CCC ATA GTT CAG C | This paper | N/A |
| **DNA plasmids** | | |
| His-CFAP418 | (1) | N/A |
| GST-CFAP418 FL | (1) | N/A |
| GST-CFAP418 CT | (1) | N/A |
| FLAG-CFAP418 | (1) | N/A |
| EGFP-CFAP418 | (1) | N/A |

| | | |
|---|---|---|
| mCherry-RAB28 | (2) | N/A |
| mCherry-RAB28 T26N | Guoxin Ying, University of Utah | N/A |
| mCherry-RAB28 Q72L | This paper | N/A |

**DATA AVAILABILITY**

The proteins pulled down in our AP-MS experiments are available in Table S1. The lipidomic, proteomic, and phosphoproteomic datasets are presented as spreadsheets in Table S2. The mass spectrometry proteomic and phosphoproteomic data have been deposited to the ProteomeXchange Consortium (http://www.proteomexchange.org) via the MASSIVE repository with the dataset identifiers MSV000089334 (P5) and MSV000089424 (P10). The mass spectrometry lipidomic data have been deposited to the Metabolomics workbench (https://www.metabolomicsworkbench.org) (3) with the study identifier ST002162. No code was generated in this study.

**METHOD DETAILS**

***Label-free quantitative MS:*** This experiment was conducted by the Mass Spectrometry Proteomics Core at Baylor College of Medicine.

MS analysis: The retinal sample preparation and high pH C18 reverse phase sample preparation were performed as described previously (4). Briefly, the tissue lysate was digested using trypsin enzyme followed by an offline high pH C18 reverse phase fractionation. The peptides were eluted in a stepwise gradient of acetonitrile (2%, 4%, 6%, 8%, 10%, 12%, 14%, 16%, 18%, 20%, 22%, 24%, 26%, 28% and 30%) and pooled into five fractions (2+12+22; 4+14+24; 6+16+26;

8+18+28; 10+20+30). These fractions were then subjected to the second dimensional low pH reversed-phase chromatography. The fractionated peptides were loaded onto a Reprosil-Pur Basic C18 (1.9 $\mu$m, Dr.Maisch GmbH, Germany) precolumn of 2 cm X 100 $\mu$m size. The precolumn was switched in-line with an in-housed 50 mm x 150 µm analytical column packed with Reprosil-Pur Basic C18 equilibrated in 0.1% formic acid/water. The peptides were analyzed using a nano-LC 1200 system (Thermo Fisher Scientific, San Jose, CA) coupled to Orbitrap Fusion™ Lumos ETD (Thermo Fisher Scientific, San Jose, CA) mass spectrometer operated in the data-dependent acquisition mode acquiring fragmentation spectra of the top 30 strongest ions and under the direct control of Xcalibur software (Thermo Scientific).

<u>Protein identification and quantification:</u> Raw files from Orbitrap Fusion Lumos were searched with Mascot algorithm (Mascot 2.4, Matrix Science) with percolator against the mouse NCBI refseq protein database (updated on 2019/01/14) in the Proteome Discoverer (PD2.1, Thermo Fisher). Dynamic modification of Oxidation, protein N-terminal Acetylation, Phosphorylation on serine, tyrosine and threonine, Deamidation on asparagine and glutamine was allowed. The precursor mass tolerance was confined within 20 ppm with fragment mass tolerance of 0.5 dalton and a maximum of two missed cleavage was allowed. The Peptide Spectrum Matches (PSMs) output from PD2.1 was grouped into gene products by 'gpGrouper' algorithm (5) and quantification was done using the label-free, intensity-based absolute quantification (iBAQ) approach and then normalized to FOT (a fraction of the total protein iBAQ amount per experiment). FOT was defined as individual protein's iBAQ divided by the total iBAQ of all identified proteins within one experiment.

***TMT-labeling quantitative MS:*** This experiment was conducted by the Thermo Fisher Center for Multiplexed Proteomics at the Harvard Medical School.

Preparation of samples for peptide isobaric labeling and MS: Retinal tissues were prepared for quantitative analysis, as previously described with some modifications (6). Tissue was lysed in 8M urea, 200 mM EPPS pH 8.0 with EDTA free cOmplete protease inhibitor cocktail and PhosSTOP phosphatase inhibitor (Roche). Total protein was quantified by micro-BCA assay (Pierce). Proteins were reduced with 5 mM TCEP and alkylated with 15 mM iodoacetamide. Total protein was precipitated with 20% trichloroacetic acid on ice, pelleted at 21,000 g, and washed twice with cold acetone. Proteins were resuspended in 200 mM EPPS, pH 8.0 and digested overnight with LysC (Wako, Japan) in a 1:50 enzyme:protein ratio followed by trypsin for 6 hours at 37°C (1:100 enzyme:protein ratio). Peptides were quantified by micro-BCA assay and 75 μg of peptide per sample were labeled with TMT10 reagents (Thermo-Fisher) for 2 hours at room temperature. Labeling reactions were quenched with 0.5% hydroxylamine and acidified with trifluoroacetic acid. Acidified peptides were combined and desalted by Sep-Pak (Waters).

Basic pH reversed-phase separation: TMT labeled peptides were solubilized in 5% acetonitrile (ACN)/10 mM ammonium bicarbonate (pH 8.0), and 300 μg of TMT labeled peptides was separated by an Agilent 300 Extend C18 column (3.5 μm particles, 4.6 mm ID and 250 mm in length). An Agilent 1260 binary pump coupled with a photodiode array detector (Thermo Scientific) was used to separate the peptides. A 45-minute linear gradient from 10% to 40% ACN in 10 mM ammonium bicarbonate pH 8.0 (flow rate of 0.6 mL/min) separated the peptide mixtures into a total of 96 fractions (36 seconds). A total of 96 fractions were consolidated into 24 samples in a checkerboard fashion, acidified with 20 μL of 10% formic acid (FA), and vacuum dried to completion. Each sample was desalted via Stage Tips and re-dissolved in 5% FA/ 5% ACN for LC-MS3 analysis.

Liquid chromatography separation and tandem mass spectrometry (LC-MS3): Data were collected using an Orbitrap Fusion mass spectrometer (Thermo Fisher Scientific) coupled with a Proxeon EASY-nLC 1200 LC pump (Thermo Fisher Scientific). Peptides were separated on a 75-µm inner diameter microcapillary column packed with 35 cm of GP-18 resin (2.6 µm, 200 Å, Sepax, Newark, DE). For each sample, ~1.5 µg of peptides were separated using a 3-hour gradient of 7–32% ACN in 0.125% FA with a flow rate of 500 nL/min.

Each analysis used an $MS^3$-based TMT method with real-time spectral searching as described previously (7). MS1 scans were acquired in the Orbitrap using a mass range of *m/z* 400 – 1500, resolution 120,000, AGC target 2.5 x $10^6$, maximum injection time 50 ms, and dynamic exclusion of 180 seconds. Data dependent top 10 $MS^2$ spectra were acquired in the ion trap with a normalized collision energy set at 35%, AGC target set to 2.5 x $10^4$ and a maximum injection time of 35 ms. MS2 spectra were searched in real time using a modified version of COMET against a mouse reference protein database (Uniprot) and common contaminants were filtered using a real-time FDR filter. Peptide spectral matches passing FDR filter and not matching to decoy peptides trigger SPS-MS3 scans collected in the Orbitrap.

Data analysis: A compendium of in-house developed software was used to convert mass spectrometric data (.RAW), file processing, monoisotopic peak correction, controlling peptide and protein level FDRs, assembling proteins from peptides, and protein quantification from peptides (8, 9). MS2 spectra were searched with COMET (10) against the mouse UniProt database (downloaded July 2014) appended with reversed protein sequences and known contaminants. COMET search parameters were 50 ppm precursor ion tolerance, 1.0005 fragment ion tolerance, fully tryptic with a maximum of 2 missed cleavage sites, Static modification of N termini and lysine residues by Ten-plex TMT tags (+ 229.162932 Da), static carboxyamidomethylation of cysteine residues (+57.02146 Da), and differential oxidation of methionine residues (+ 15.99492 Da). Peptide and protein were filtered to an FDR of less than 1% by applying the target-decoy database search and linear discriminant analysis (8, 9).

Proteins were quantified as previously described (11). A 0.003 *m/z* window centered on the theoretical m/z value of each of the six reporter ions and the intensity of the signal closest to the theoretical m/z value was recorded. Reporter ion intensities were adjusted based on the overlap of isotopic envelopes of all reporter ions (as determined by the manufacturer). Only peptides with a total summed reporter ion signal to noise greater than 100 and MS2 isolation specificity greater than 0.5 were used for protein quantification.

***<u>Lipidomic MS</u>:*** This experiment was conducted by the Metabolomics Core at the University of Utah.

<u>Chemicals:</u> LC-MS-grade solvents and mobile phase modifiers were obtained from Honeywell Burdick & Jackson, Morristown, NJ (ACN, isopropanol, FA), Fisher Scientific, Waltham, MA (methyl *tert*-butyl ether) and Sigma–Aldrich/Fluka, St. Louis, MO (ammonium formate).

<u>Sample preparation:</u> Extraction of lipids was carried out using a biphasic solvent system of cold methanol, methyl *tert*-butyl ether (MTBE), and water with some modifications (12). In a randomized sequence, tissue lipids were extracted in bead-mill tubes (ceramic 1.4 mm, Mo-Bio, Qiagen, Germantown, MD) containing a solution of 225 μL methanol, 750 μL MTBE, and internal standards (Lipid standard Mouse SPLASH LipidoMix at 10 μL per sample, Avanti Polar

Lipids, Alabaster, AL). Samples were homogenized in one 30-second cycle and rested on ice for 1 hour with occasional vortexing. Then, 188 µL of PBS was added followed by a brief vortex. Samples were centrifuged at 14,000 x *g* for 10 minutes at 4 °C, and the upper phases were collected. Another aliquot of 750 µL MTBE was added to the bottom aqueous layer followed by a brief vortex. Samples were then centrifuged at 14,000 x *g* for 10 minutes at 4 °C, the upper phases were combined and evaporated to dryness under speedvac. Lipid extracts were reconstituted in 250 µL of mobile phase B and transferred to an LC-MS vial for analysis. Concurrently, a process blank sample was prepared and then a pooled quality control (QC) sample was prepared by taking equal volumes (~50 µL) from each sample after final resuspension.

LC-MS analysis: Lipid extracts were separated on an Acquity UPLC CSH C18 column (2.1 x 100 mm; 1.7 µm) coupled to an Acquity UPLC CSH C18 VanGuard precolumn (5 × 2.1 mm; 1.7 µm) (Waters, Milford, MA) maintained at 65 °C connected to an Agilent HiP 1290 Sampler, Agilent 1290 Infinity pump, and Agilent 6545 Accurate Mass Q-TOF dual AJS-ESI mass spectrometer (Agilent Technologies, Santa Clara, CA). Samples were analyzed in a randomized order in both positive and negative ionization modes in separate experiments acquiring with the scan range m/z 100 – 1700. For positive mode, the source gas temperature was set to 225 °C, with a drying gas flow of 11 L/minute, nebulizer pressure of 40 psig, sheath gas temp of 350 °C and sheath gas flow of 11 L/minute. VCap voltage is set at 3500 V, nozzle voltage 500V, fragmentor at 110 V, skimmer at 85 V and octopole RF peak at 750 V. For negative mode, the source gas temperature was set to 300 °C, with a drying gas flow of 11 L/minute, a nebulizer pressure of 30 psig, sheath gas temp of 350 °C and sheath gas flow 11 L/minute. VCap voltage was set at 3500 V, nozzle voltage 75 V, fragmentor at 175 V, skimmer at 75 V and octopole RF peak at 750 V. Mobile phase A consisted of ACN:$H_2O$ (60:40 *v/v*) in 10 mM ammonium formate and 0.1% FA, and mobile phase B consisted of IPA:ACN:$H_2O$ (90:9:1 *v/v/v*) in 10 mM ammonium formate and 0.1% FA. The chromatography gradient for both positive and negative modes started at 15% mobile phase B then increased to 30% B over 2.4 min. It sequentially increased to 48% B from 2.4 – 3.0 min, 82% B from 3 – 13.2 min, and 99% B from 13.2 – 13.8 min where it's held until 16.7 min and returned to the initial conditions and equilibrated for 5 min. Flow was 0.4 mL/minute throughout, with injection volumes of 2 µL for positive and 10 µL for negative mode, and iterative, tandem mass spectrometry was conducted using the same LC gradient at collision energies of 20 V and 27.5 V, respectively.

<u>LC-MS data processing:</u> For data processing, Agilent MassHunter (MH) Workstation and software packages MH Qualitiative and MH Quantitative were used. The pooled QC (n=8) and process blank (n=4) were injected throughout the sample queue to ensure the reliability of acquired lipidomics data. For lipid annotation, accurate mass and MS/MS matching was used with the Agilent Lipid Annotator library. Results from the positive and negative ionization modes from Lipid Annotator were merged based on the class of lipid identified. Data exported from MH Quantitative was evaluated using Excel where initial lipid targets were parsed based on the following criteria. Only lipids with relative standard deviations less than 30% in QC samples were used for data analysis. Additionally, only lipids with background AUC counts in process blanks that were less than 30% of QC were used for data analysis. The parsed excel data tables were normalized based on the ratio to class-specific internal standards, then to sum prior to statistical analysis.

***<u>Affinity-purification coupled with mass spectrometry:</u>***

Affinity purification: Mouse and bovine retinas were homogenized and incubated in lysis buffer (200 mM NaCl, 20 mM Tris, 0.5% Triton X-100, 1 mM TCEP, protease inhibitor cocktail, pH8.0) for 30 minutes at 4°C. The homogenates were then centrifuged at 100,000 x g at 4°C for 30 minutes. For mouse retinas, the affinity purification was conducted by incubating the obtained supernatants with our custom-made CFAP418 antibody and subsequently protein G sepharose (Fisher scientific, Waltham, MA) for 30 minutes each or with the CFAP418 antibody cross-linked with protein G for 30 minutes total. The protein G beads were spun down at 3,500 x g for 30 seconds and washed four times with the lysis buffer. For bovine retinas, the supernatants were incubated with recombinant GST-tagged CFAP418 proteins expressed in One Shot™ BL21 Star™ (DE3) cells (ThermoFisher, Waltham, MA) and purified using Pierce™ Glutathione Superflow Agarose (ThermoFisher, Waltham, MA). The precipitated proteins were eluted from the protein G sepharose or Glutathione agarose beads using 2X SDS sample loading buffer and run on a 10% SDS-PAGE. Each protein lane was cut into 2-4 sections and submitted to the Taplin Mass Spectrometry Facility at Harvard Medical School for protein identification. More details are shown in Figure 1B.

Gel band processing: Gel bands were cut into approximately 1-$mm^3$ pieces and subjected to a modified in-gel trypsin digestion procedure (13). Briefly, gel pieces were washed and dehydrated with ACN for 10 minutes. After ACN removal, pieces were completely dried in a speed-vac and rehydrated at 4ºC for 45 minutes with 50 mM ammonium bicarbonate solution containing 12.5 ng/µl modified sequencing-grade trypsin (Promega, Madison, WI). After removal of excess trypsin, replacement with 50 mM ammonium bicarbonate solution, and incubation at 37ºC overnight, peptides were extracted by removing the ammonium bicarbonate solution, one wash with a solution containing 50% ACN and 1% FA, and dried in a speed-vac for ~1 hour.

LC-MS/MS analysis: Gel extraction samples were reconstituted in 5 - 10 µl of HPLC solvent A (2.5% ACN and 0.1% FA). A reverse-phase HPLC capillary column was generated by packing 2.6 µm-C18 spherical silica beads into a fused silica capillary (100 µm inner diameter x ~30 cm length) (14). After equilibrating the column, samples were loaded through a Famos auto sampler (LC Packings, San Francisco CA). Peptides were eluted with increasing concentrations of solvent B (97.5% ACN and 0.1% FA). As peptides were eluted, they were subjected to electrospray ionization and entered into an LTQ Orbitrap Velos Pro ion-trap mass spectrometer (ThermoFisher Scientific, Waltham, MA). Peptides were detected, isolated, and fragmented to produce a tandem mass spectrum of specific fragment ions. Peptide sequences and protein identity were determined by matching protein databases with the acquired fragmentation pattern by Sequest (ThermoFisher Scientific, Waltham, MA) (15). All databases included a reversed version of all the sequences and the data were filtered between a 1 and 2% peptide FDR.

**Supplemental Information**

**Table S1: Proteins immunoprecipitated by CFAP418 antibody from mouse retinas and proteins pulled down by GST-CFAP418 baits from bovine retinas.**

**Table S2: Original data from quantitative proteomic, phosphoproteomic, and lipidomic studies.** Sheet 1: Proteomic data at P5. Sheet 2: Phosphoproteomic data at P5. Sheet 3: Proteomic data at P10. Sheet 4: Phosphoproteomic data at P10. Sheet 5: lipidomic data at P10.

**Table S3: DE proteins, DP proteins, and altered lipid species identified in *Cfap418*$^{-/-}$ retinas by MetaboAnalyst.** Reduced DE and DP proteins and lipids are highlighted in blue. Increased DE and DP proteins and lipids are highlighted in yellow. Other proteins and lipids have a *p*-value or an adjusted *p*-value between 0.05 and 0.1. Sheet 1: Proteomic data at P5. Sheet 2: Phosphoproteomic data at P5. Sheet 3: Proteomic data at P10. Sheet 4: Phosphoproteomic data at P10. Sheet 5: lipidomic data at P10.

**Table S4: GSEA reports of biological pathways, molecular functions, and cellular compartments that are affected by *Cfap418* knockout at P5** (sheet 1: down-regulated; sheet 2: up-regulated) **and P10** (sheet 3: down-regulated; sheet 4: up-regulated)**.**

**Table S5: DE and DP proteins identified in *Cfap418*$^{-/-}$ retinas at both P5 and P10.** Sheet 1: the proteomic and phosphoproteomic data. Sheet 2: Known information and references of the shared DE proteins.

### Supplemental Figure Legends

**Figure S1: CFAP418 protein sequence alignment and evaluation of OS membrane protein synthesis, degradation, and ER stress in *Cfap418*$^{-/-}$ retinas.** (**A**) CFAP418 sequence alignment across different species. Red, blue, and black residues represent highly, mildly, and not conserved residues, respectively. Patient mutations are shown on the top of sequences. Splice site, nonsense, and frameshift mutations are denoted by vertical lines, and missense mutations are highlighted in grey. **(B)** Pulse labeling using [$^{35}$S] methionine for up to 2 hours shows no obvious reduction in newly synthesized rhodopsin (RHO) in *Cfap418*$^{-/-}$ retinas at P12, compared with *Cfap418*$^{+/-}$ littermate retinas. DPYSL2 is a cytoplasmic protein and was used as a negative control. Left, autoradiograms of pulse-labeled and immunoprecipitated RHO and DPYSL2 proteins on SDS-PAGE. Right, quantification of the RHO and DPYSL2 radioactivity from the autoradiograms. (**C**) Chase labeling of *Cfap418*$^{+/-}$ and *Cfap418*$^{-/-}$ littermate retinal explants for 3 hours at P12 and overnight at P8 using [$^{35}$S] methionine shows no obvious changes in the degradation of RHO or transducin $\alpha$ subunit (GNAT1). Left, representative autoradiograms of chase-labeled and immunoprecipitated RHO and GNAT1 proteins on SDS-PAGE. Right, quantitation of radioactive RHO and GNAT1 signals. (**D**) RT-qPCR analysis shows no changes in the PERK, IRE1$\alpha$, and ATF6 mRNAs in *Cfap418*$^{-/-}$ retinas at P15 and P30. Each dot represents the retinas of an individual mouse. Mean ± SEM is shown. (**E**) Immunoblot analysis demonstrates no cleavage of ATF6 into a 50-kDa fragment or increase of IRE1$\alpha$ phosphorylation in *Cfap418*$^{-/-}$ retinas at P16 and P30. $\gamma$-tubulin is a sample loading control.

**Figure S2: Binding of CFAP418 to lipids and lipid changes in *Cfap418*$^{-/-}$ retinas.** **(A)** Both His-tagged and GST-tagged mouse CFAP418 proteins bind to phosphatidic acid (PA) on membrane lipid strips. His-CFAP418 also binds to lysophosphatidic acid (LPA) weakly. The lipid arrangement is the same on the two strips. **(B)** Heatmaps show the abundances of individual lipid species in each sample before and after the elimination of the outlier, sample 14. **(C)** Pareto abundance histograms of various lipid categories in *Cfap418*$^{+/-}$ and *Cfap418*$^{-/-}$ retinas at P10. Cumulative lines are shown. The numbers on top of each bin indicate the numbers of lipid species in the bin. Red and black underlines indicate the increased and reduced lipid categories in *Cfap418*$^{-/-}$ retinas. **(D)** Pareto abundance histograms of various acyl chains in *Cfap418*$^{+/-}$ and *Cfap418*$^{-/-}$ retinal membrane lipids at P10. Legends are the same as in **C**. **(E)** The seven PC species that are altered in P10 *Cfap418*$^{-/-}$ retinas. Data from individual mice and mean ± SEM are shown. * and **: $p < 0.05$ and 0.01, respectively, based on Student's t-test on the MetaboAnalyst processed data.

**Figure S3: Colocalization analysis of CFAP418 with ESCRT and RAB28 proteins. (A)** Representative COS-7 cells transfected with FLAG-CFAP418 and double-immunostained for FLAG and ESCRT proteins. **(B)** Representative COS-7 cells double-transfected with FLAG-CFAP418 and mCherry-RAB28 mutant proteins. Cells were immunostained using FLAG and mCherry antibodies. Arrows point to the abnormally accumulated vacuoles. Framed regions are amplified and shown on the right with merged and individual channels. PCCs are indicated as mean ± SEM. n, number of cells analyzed. Scale bars: 10 μm.

**Figure S4: Localization of vesicular trafficking proteins and SLC1A1 in *Cfap418*$^{-/-}$ photoreceptors. (A)** Double staining of RAB28 and rhodopsin in *Cfap418*$^{+/-}$ and *Cfap418*$^{-/-}$ photoreceptors at P10. As reported previously (2), RAB28 is in photoreceptor OS, IS, and OPL.

The strongest RAB28 signal is present in the OS, marked by rhodopsin. There is no significant difference in RAB28 distribution between P10 *Cfap418*$^{+/-}$ and *Cfap418*$^{-/-}$ photoreceptors. **(B)** SLC1A1 is present in all photoreceptor layers at P10. SV2 labels the photoreceptor OPL. There is no significant difference in SLC1A1 distribution between P10 *Cfap418*$^{+/-}$ and *Cfap418*$^{-/-}$ photoreceptors. **(C)** TFG and VPS4B proteins are localized normally in *Cfap418*$^{-/-}$ photoreceptors, compared with littermate *Cfap418*$^{+/-}$ photoreceptors at P10. **(D)** Early endosomal proteins EEA1 and RAB5, late endosomal protein RAB7, and recycling endosomal protein RAB11 are localized normally in *Cfap418*$^{-/-}$ photoreceptors, compared with littermate *Cfap418*$^{+/-}$ photoreceptors at P21. Scale bars: 10 μm.

**Figure S5: Distribution of PRKCA and phosphorylated PRKCA in developing and mature mouse retinas.** Pan- and pT497-PRKCA immunoreactivities are present throughout the retina including various photoreceptor layers at P5 (left) and are mainly located in rod bipolar cells in the retina at P21 (right). Scale bars, 10 μm.

**Figure S6: Expressions of IRD-associated proteins in *Cfap418*$^{-/-}$ retinas.** The expressions of proteins encoded by known retinal disease genes (RetNet) and detected in our P5 and P10 quantitative proteomic studies are shown. Totally, 25 proteins are differentially expressed in *Cfap418*$^{-/-}$ retinas. Among them, proteins in the BBSome, OS, and spliceosome tend to be more affected. Note that the clustering of proteins in this figure is based on the current knowledge of functional and direct protein associations. The associations of CFAP418 with BBSome and RAB28 revealed in this study are novel and not shown. The gradient from dark blue to dark red reduced to the most increased fold change in *Cfap418*$^{-/-}$ retinas. Dashed border and square-shaped node denote the statistical significance ($p < 0.05$) at P5 and P10, respectively. The red color of node labels depicts all the DE proteins found in P5 and P10 *Cfap418*$^{-/-}$ retinas. The cellular functions and compartments of protein clusters circled by red lines are labeled.

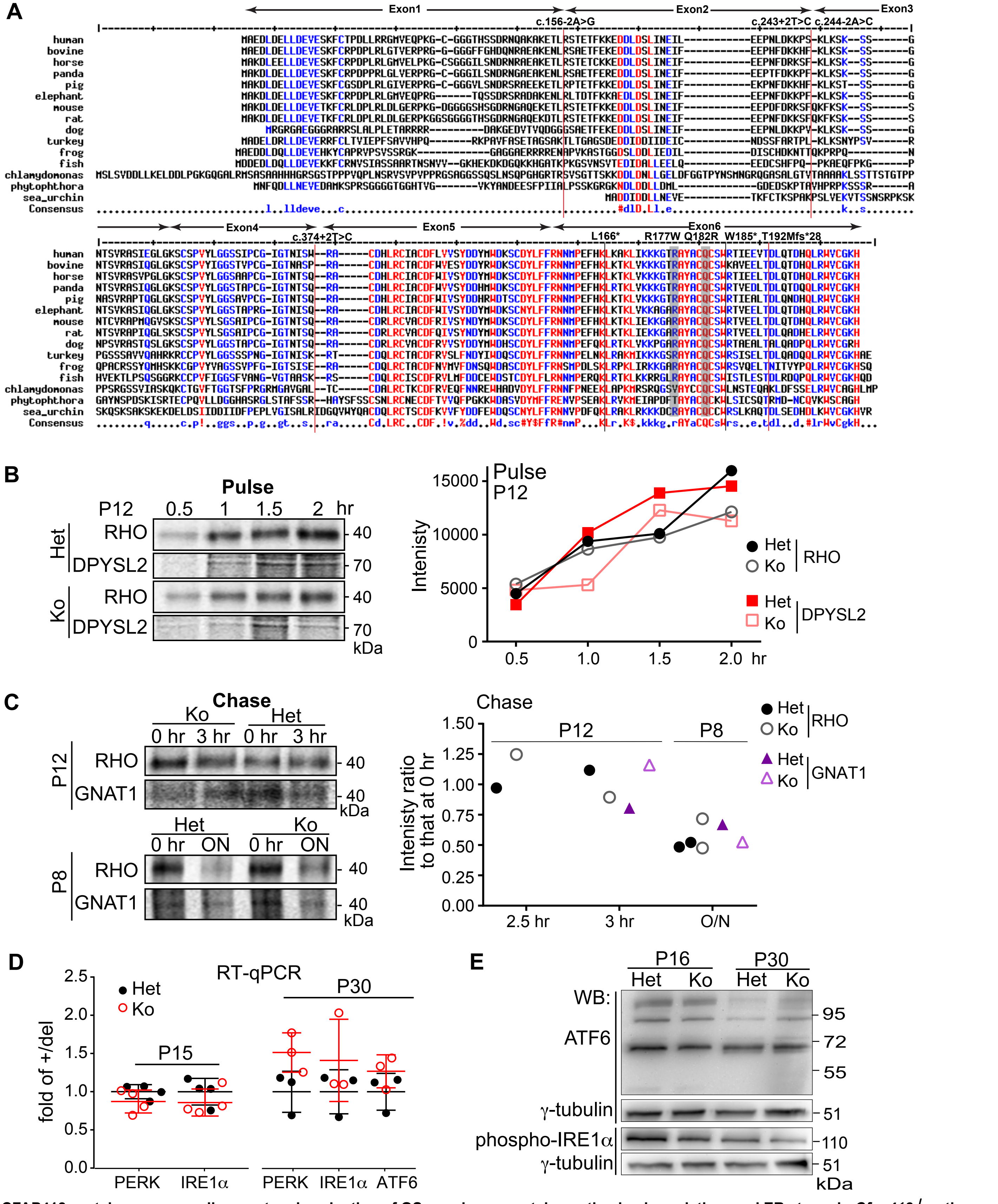


**Figure S1: CFAP418 protein sequence alignment and evaluation of OS membrane protein synthesis, degradation, and ER stress in *Cfap418*$^{-/-}$ retinas.** (**A**) CFAP418 sequence alignment across different species. Red, blue, and black residues represent highly, mildly, and not conserved residues, respectively. Patient mutations are shown on the top of sequences. Splice site, nonsense, and frameshift mutations are denoted by vertical lines, and missense mutations are highlighted in grey. (**B**) Pulse labeling using [$^{35}$S] methionine for up to 2 hours shows no obvious reduction in newly synthesized rhodopsin (RHO) in *Cfap418*$^{-/-}$ retinas at P12, compared with *Cfap418*$^{+/-}$ littermate retinas. DPYSL2 is a cytoplasmic protein and was used as a negative control. Left, autoradiograms of pulse-labeled and immunoprecipitated RHO and DPYSL2 proteins on SDS-PAGE. Right, quantification of the RHO and DPYSL2 radioactivity from the autoradiograms. (**C**) Chase labeling of *Cfap418*$^{+/-}$ and *Cfap418*$^{-/-}$ littermate retinal explants for 3 hours at P12 and overnight at P8 using [$^{35}$S] methionine shows no obvious changes in the degradation of RHO or transducin subunit (GNAT1). Left, representative autoradiograms of chase-labeled and immunoprecipitated RHO and GNAT1 proteins on SDS-PAGE. Right, quantitation of radioactive RHO and GNAT1 signals. (**D**) RT-qPCR analysis shows no changes in the PERK, IRE1α, and ATF6 mRNAs in *Cfap418*$^{-/-}$ retinas at P15 and P30. Each dot represents the retinas of an individual mouse. Mean ± SEM is shown. (**E**) Immunoblot analysis demonstrates no cleavage of ATF6 into a 50-kDa fragment or increase of IRE1α phosphorylation in *Cfap418*$^{-/-}$ retinas at P16 and P30. γ-tubulin is a sample loading control.

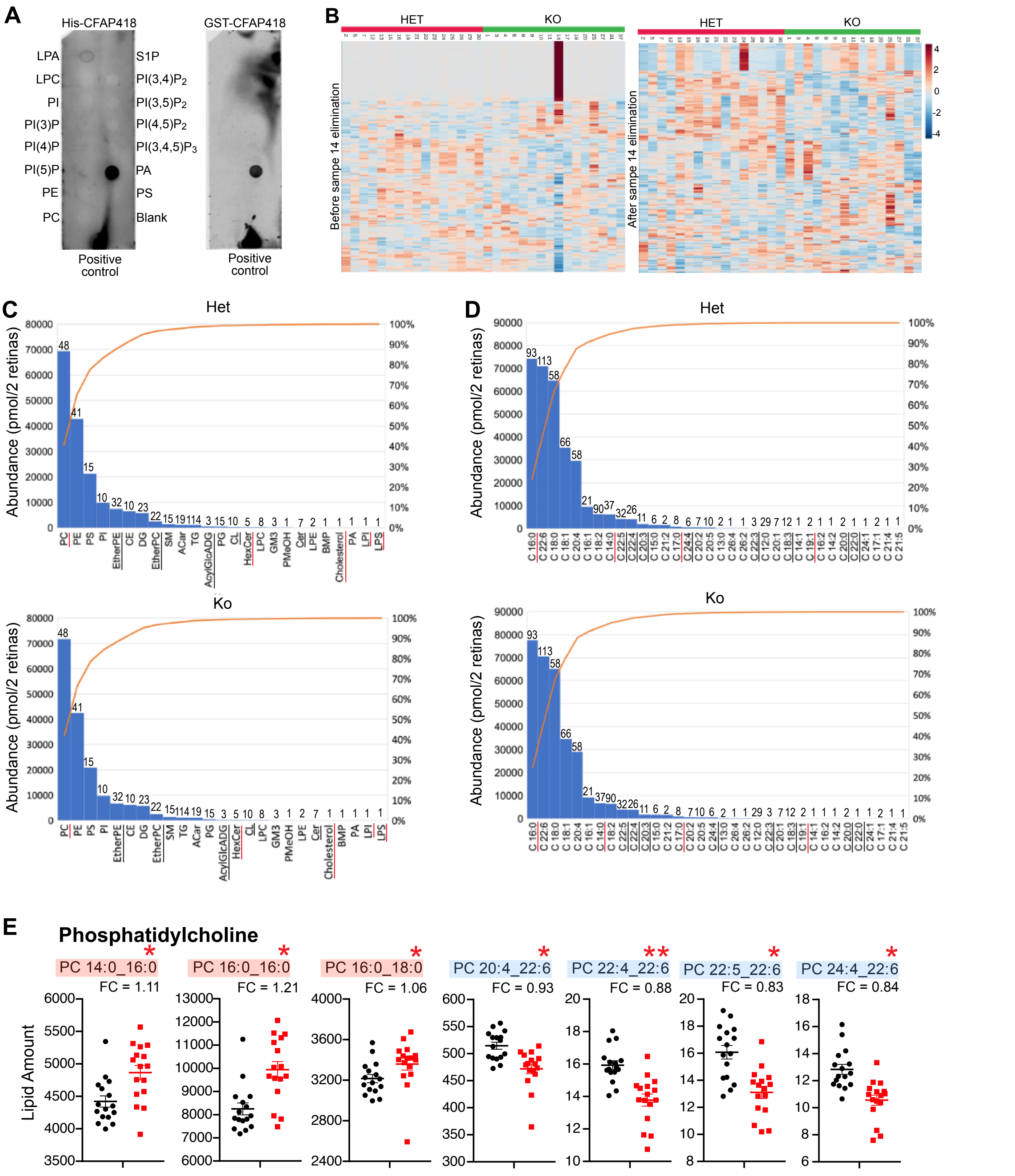


**Figure S2: Binding of CFAP418 to lipids and lipid changes in *Cfap418*$^{-/-}$ retinas.** (**A**) Both His-tagged and GST-tagged mouse CFAP418 proteins bind to phosphatidic acid (PA) on membrane lipid strips. His-CFAP418 also binds to lysophosphatidic acid (LPA) weakly. The lipid arrangement is the same on the two strips. (**B**) Heatmaps show the abundances of individual lipid species in each sample before and after the elimination of the outlier, sample 14. (**C**) Pareto abundance histograms of various lipid categories in *Cfap418*$^{+/-}$ and *Cfap418*$^{-/-}$ retinas at P10. Cumulative lines are shown. The numbers on top of each bin indicate the numbers of lipid species in the bin. Red and black underlines indicate the increased and reduced lipid categories in *Cfap418*$^{-/-}$ retinas. (**D**) Pareto abundance histograms of various acyl chains in *Cfap418*$^{+/-}$ and *Cfap418*$^{-/-}$ retinal membrane lipids at P10. Legends are the same as in **C**. (**E**) The seven PC species that are altered in P10 *Cfap418*$^{-/-}$ retinas. Data from individual mice and mean ± SEM are shown. * and **: $p < 0.05$ and 0.01, respectively, based on Student's t-test on the MetaboAnalyst processed data.

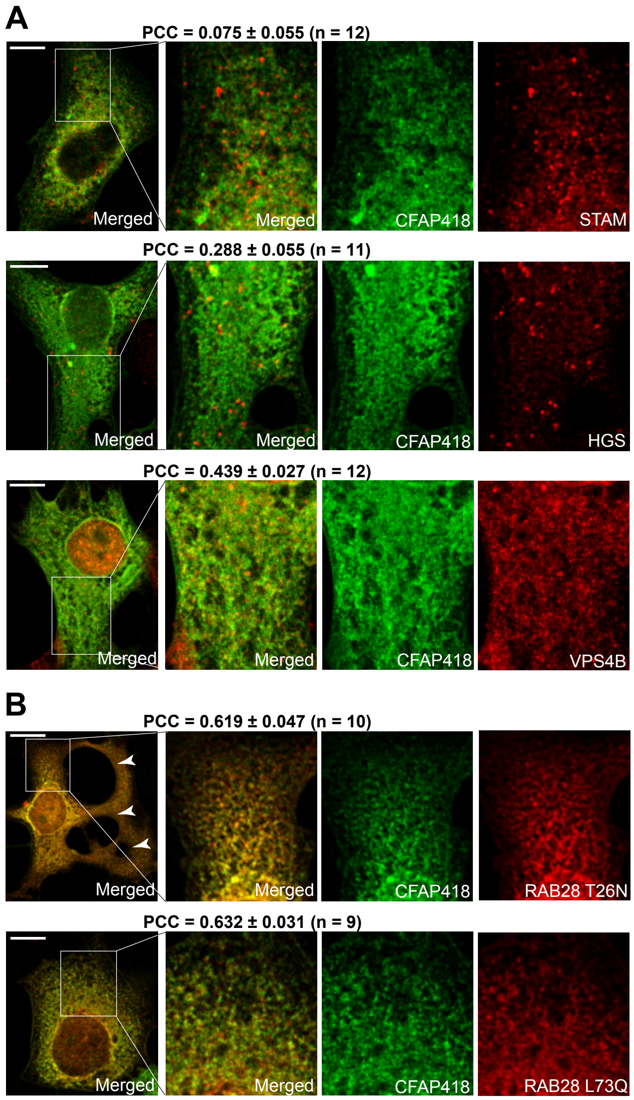


**Figure S3: Colocalization analysis of CFAP418 with ESCRT and RAB28 proteins.** (**A**) Representative COS-7 cells transfected with FLAG-CFAP418 and double-immunostained for FLAG and ESCRT proteins. (**B**) Representative COS-7 cells double-transfected with FLAG-CFAP418 and mCherry-RAB28 mutant proteins. Cells were immunostained using FLAG and mCherry antibodies. Arrows point to the abnormally accumulated vacuoles. Framed regions are amplified and shown on the right with merged and individual channels. PCCs are indicated as mean ± SEM. n, number of cells analyzed. Scale bars: 10 μm.

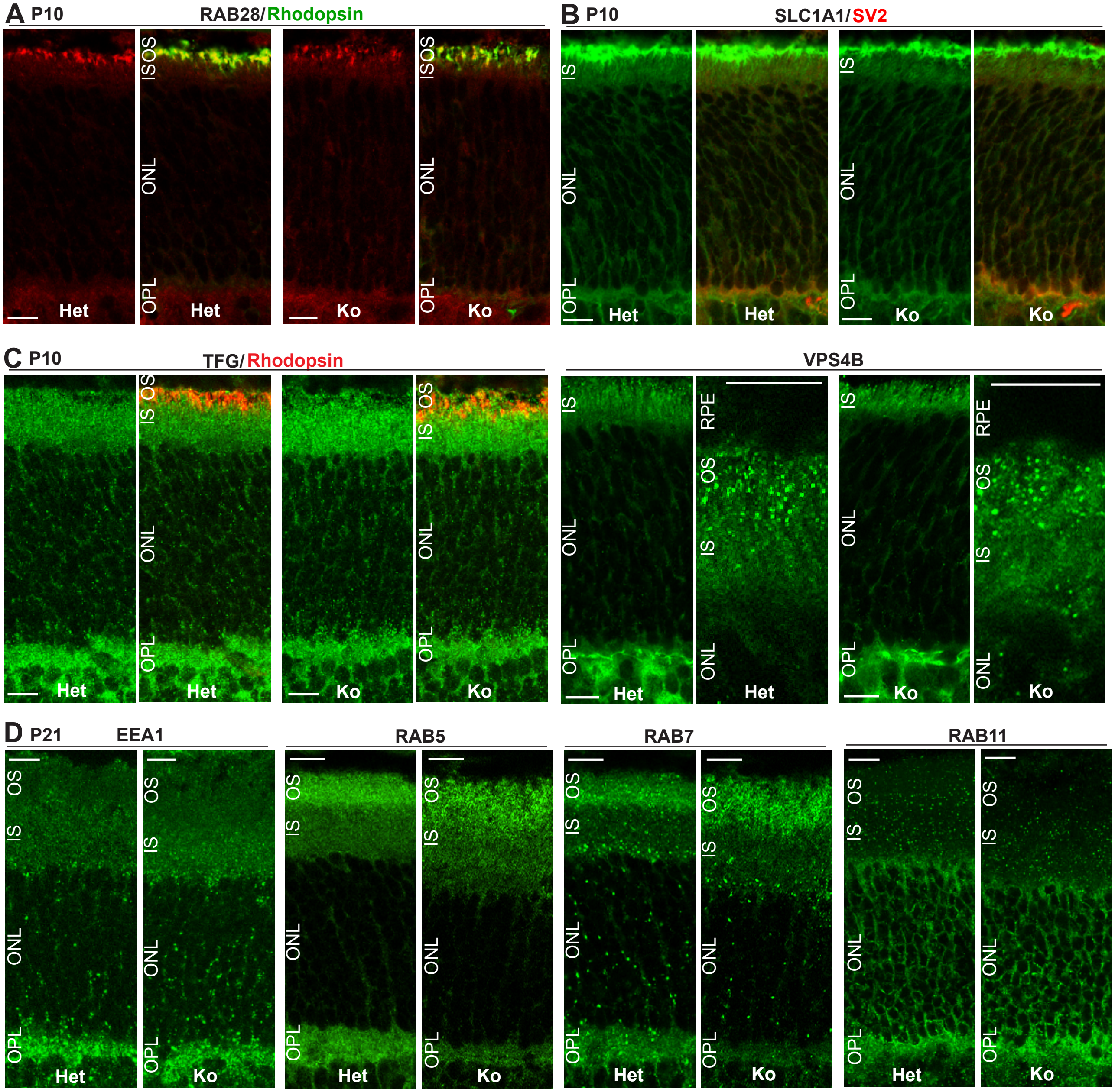


**Figure S4: Localization of vesicular trafficking proteins and SLC1A1 in *Cfap418*$^{-/-}$ photoreceptors.** (**A**) Double staining of RAB28 and rhodopsin in *Cfap418*$^{+/-}$ and *Cfap418*$^{-/-}$ photoreceptors at P10. As reported previously (22), RAB28 is in photoreceptor OS, IS, and OPL. The strongest RAB28 signal is present in the OS, marked by rhodopsin. There is no significant difference in RAB28 distribution between P10 *Cfap418*$^{+/-}$ and *Cfap418*$^{-/-}$ photoreceptors. (**B**) SLC1A1 is present in all photoreceptor layers at P10. SV2 labels the photoreceptor OPL. There is no significant difference in SLC1A1 distribution between P10 *Cfap418*$^{+/-}$ and *Cfap418*$^{-/-}$ photoreceptors. (**C**) TFG and VPS4B proteins are localized normally in *Cfap418*$^{-/-}$ photoreceptors, compared with littermate *Cfap418*$^{+/-}$ photoreceptors at P10. (**D**) Early endosomal proteins EEA1 and RAB5, late endosomal protein RAB7, and recycling endosomal protein RAB11 are localized normally in *Cfap418*$^{-/-}$ photoreceptors, compared with littermate *Cfap418*$^{+/-}$ photoreceptors at P21. Scale bars: 10 μm.

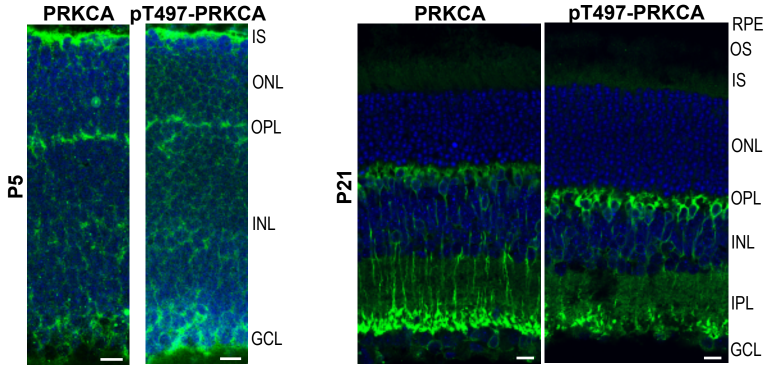


**Figure S5: Distribution of PRKCA and phosphorylated PRKCA in developing and mature mouse retinas.** Pan- and pT497-PRKCA immunoreactivities are present throughout the retina including various photoreceptor layers at P5 (left) and are mainly located in rod bipolar cells in the retina at P21 (right). Scale bars, 10 μm.

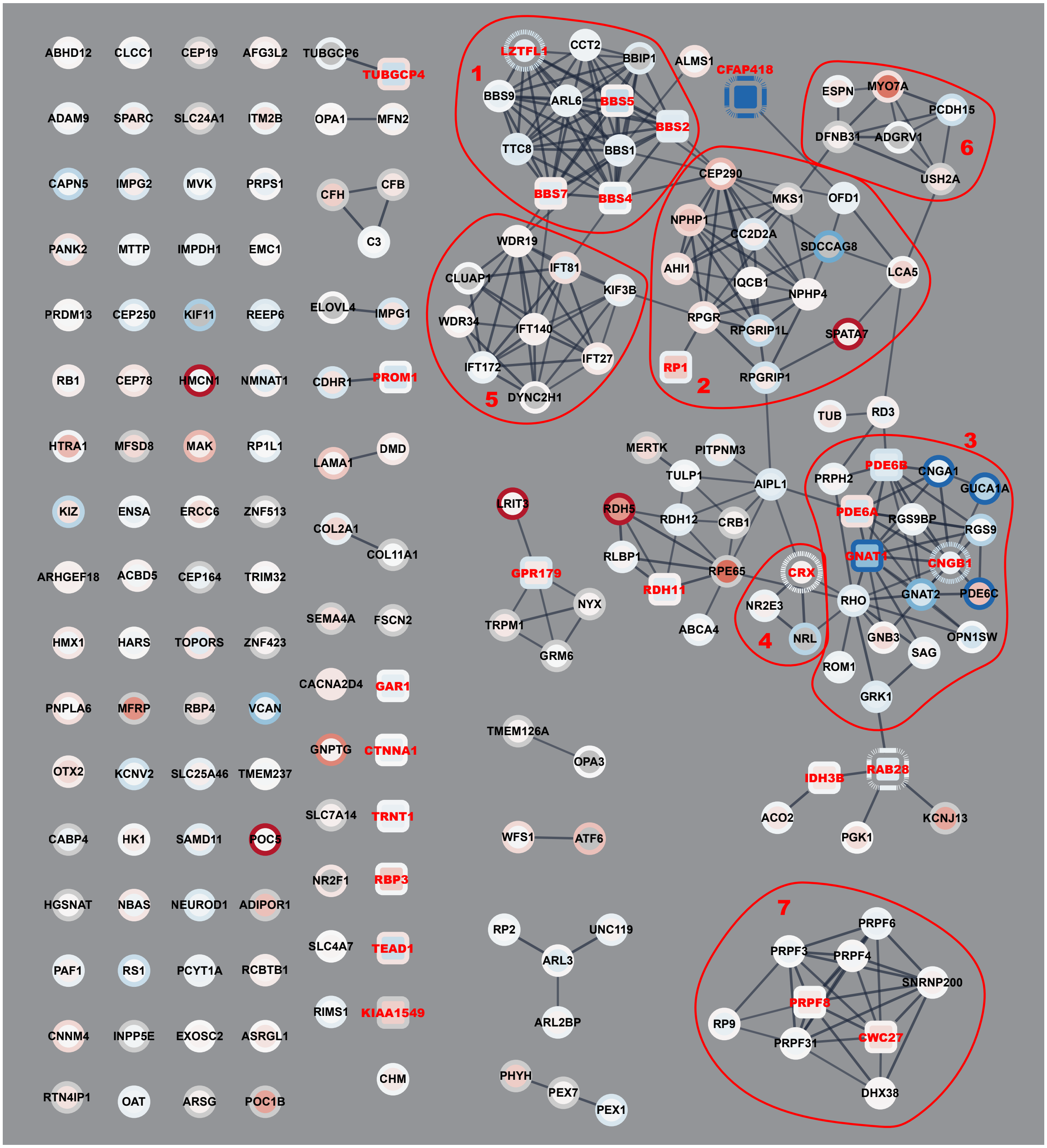


1. BBSome proteins
2. Connecting cilium proteins
3. Outer segment proteins
4. Transcription factors
5. Intraflagellar transport proteins
6. Usher proteins
7. Spliceosome proteins

**Figure S6: Expressions of IRD-associated proteins in *Cfap418*$^{-/-}$ retinas.** The expressions of proteins encoded by known retinal disease genes (RetNet) and detected in our P5 and P10 quantitative proteomic studies are shown. Totally, 25 proteins are differentially expressed in *Cfap418*$^{-/-}$ retinas. Among them, proteins in the BBSome, OS, and spliceosome tend to be more affected. Note that the clustering of proteins in this figure is based on the current knowledge of functional and direct protein associations. The associations of CFAP418 with BBSome and RAB28 revealed in this study are novel and not shown. The gradient from dark blue to dark red of nodes and node borders (P5 and P10, respectively) indicates the gradient from the most reduced to the most increased fold change in *Cfap418*$^{-/-}$ retinas. Dashed border and square-shaped node denote the statistical significance ($p < 0.05$) at P5 and P10, respectively. The red color of node labels depicts all the DE proteins found in P5 and P10 *Cfap418*$^{-/-}$ retinas. The cellular functions and compartments of protein clusters circled by red lines are labeled.
